# MIND the gap: methodological considerations and guidance for structural MRI similarity network analysis with MIND

**DOI:** 10.64898/2026.08.28.747811

**Authors:** A. Davies, E. D. Hutchings, B. Chidiac, M. H. Garvey, S. K. Crockford, I. Sebenius, R. A. I. Bethlehem, E. T. Bullmore, S. E. Morgan

## Abstract

Structural similarity networks quantify the similarity of structural properties across cortical regions, providing a macroscopic window onto the organisation of cortical architecture^1–3^. Morphometric inverse divergence (MIND) is a multivariate metric of similarity between cortical areas, based on the Kullback-Leibler (KL) divergence between areal distributions of multiple MRI features or morphometric variables locally measured at voxel or vertex resolution. MIND has demonstrated technical robustness and biological validity^4^ and is increasingly widely used as a measure of cortico-cortical similarity in clinical and developmental network neuroscience^2^. Here we provide in-depth methodological background on KL divergence and MIND, highlighting possible sources of bias, critical user decision points in the design of a MIND processing pipeline, and recommendations for technical risk mitigation in using MIND as a metric of cortical similarity. We use simulated data and observational MRI datasets from adults (UK Biobank, N = 500 T1-weighted and diffusion scans) and neonates (Developing Human Connectome Project, N = 752 T2-weighted scans), to show how the estimator of KL divergence implemented in MIND is potentially influenced or biased by five properties of input MRI feature maps: (i) their smoothness; (ii) the proportion of identical values; (iii) analysis in native or common space and the choice of vertex mesh resolution; (iv) parcellation choice; and (v) covariance between input features. We offer principled and practical guidance for investigators wanting to specify and implement the MIND processing pipeline that is best suited to the constraints and opportunities of the MRI data available to them. These recommendations outline which pipeline steps should be used sparingly, such as vertex map smoothing; which should be used with informed caution, such as parcellation choice or vertex mesh resampling; and which could be newly implemented for more robust estimation of MIND, such as the use of principal component analysis to preprocess multivariate MRI features. To support further development of structural MRI similarity network analysis, and wider adoption of robust MIND methods, we also publish the code used to generate the results in this paper as an open resource.

## 1 Introduction

Structural MRI similarity analysis has emerged as a useful framework to measure coordinated cortical patterning of multiple geometric and tissue-composition features across areas of the human cortical sheet in an individual brain scan^1,3^. Conceptually, structural similarity networks are a macro-scale network representation of the cortical ‘architectome’^2^, akin to the seminal micro-scale maps of cytoarchitectonic variation represented by Brodmann’s discrete areal parcellation^5^ or von Economo’s continuous trends of cortical differentiation^6^. Methodologically, there are several options available for estimating within-subject structural similarity^1,3^, for example based on correlation^7,8^, or information theoretic measures of distributional divergence such as the Kullback-Leibler (KL) divergence^4,9,10^. As these approaches become more widely used for clinical and developmental studies of human cortical anatomy, it is timely to review their technical foundations and potential biases or limitations in support of their well-informed implementation and interpretation.

Morphometric inverse divergence (MIND) estimates pairwise inter-areal similarity using the inverse of the KL divergence between multivariate distributions of morphometric MRI features measured in each cortical area^4^. By MRI feature, we mean any macrostructural or microstructural variable that can be locally measured at vertex or voxel resolution throughout the cortex. Highly divergent distributions, representing cortical areas that are morphologically distinct from each other, have low pairwise MIND values, while highly similar distributions, representing cortical areas that are morphologically convergent with each other, have high pairwise MIND values. Relative to prior methods of morphometric similarity analysis based on estimating the inter-areal correlation between regional mean MRI feature vectors^7^, MIND has enhanced performance on a roster of technical and biological validation criteria^4,11,12^.

MIND similarity between cortical areas has been robustly triangulated with biological benchmarks in humans^4^, non-human primates^13^ and rats^14^. As expected, given prior links between inter-areal microstructural similarity and axonal connectivity^15,16^, MIND was positively correlated with connectivity weights between cortical areas in primate and rodent tract-tracing data. In all examined species, MIND was correlated with cytoarchitectonic data on inter-areal similarity at microscopic scale, and single-cell or whole genome transcriptional data on inter-areal gene co-expression. MIND phenotypes were heritable in humans, with an underlying genetic architecture decomposable into two major gradients that spatially aligned with previously established phylogenetic axes of cortical differentiation originating from two discrete loci of primordial cortex^17,18^.

On the basis of these technical and biological validations, MIND has been applied increasingly and at scale, including to studies of normative brain development and aging in humans^19^ and other species^13,14,20^ as well as clinical case-control studies of neuropsychiatric^21–26^ and neurodegenerative disorders^27,28^. Collectively, these studies indicate that MIND can be adaptive and informative across a range of available MRI datasets and experimental designs for clinical and developmental neuroscience. However, wider use of MIND has naturally also raised methodological questions regarding how to optimise estimation pipelines for robustness against known potential biases, and how to customise MRI feature selection from the large array of macrostructural and microstructural morphometric variables that are locally measurable by contemporary multi-parameter MRI sequences^29,30^.

To address these questions, we start with a technical primer on how MIND is estimated, highlighting the central role of KL divergence and the theoretical vulnerabilities to bias in KL estimation from multivariate areal distributions of morphometric MRI features. On this foundation, we then report our two-pronged approach to methodological validation of MIND, using both simulated data and experimental MRI data (**Figure 1**).

**Figure 1:**
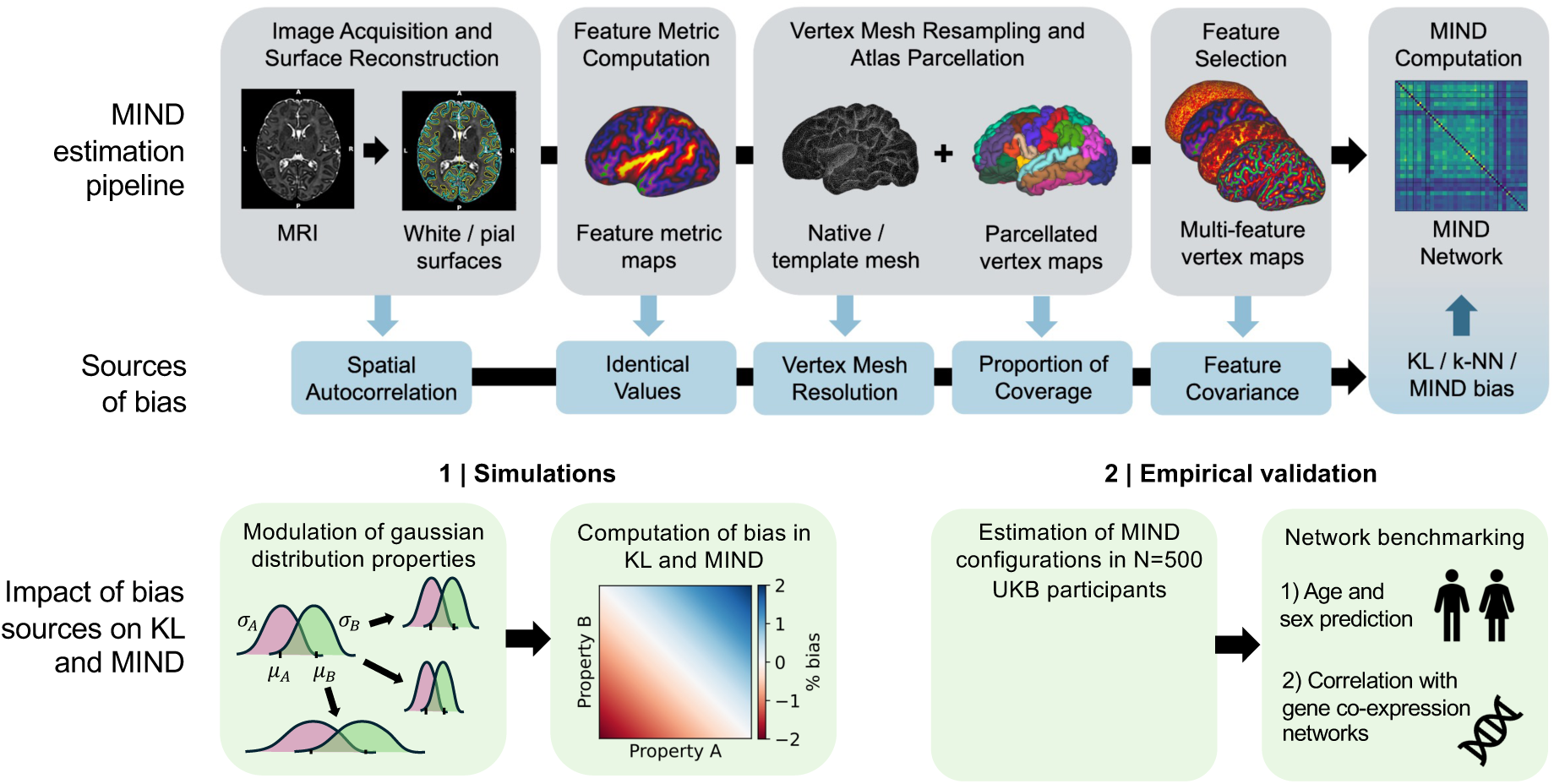
Study overview. Top row: a typical MIND estimation pipeline includes volumetric MRI preprocessing, surface interpolation, computation of feature metrics at a specified vertex mesh resolution, parcellation, feature selection, and MIND estimation. Middle row: bias can be introduced in each of these preprocessing stages. Bottom row: our approach to evaluating the effects of bias on KL divergence and MIND involved: 1) computing bias in KL and MIND by modulating distributional properties of simulated Gaussians, estimating KL and MIND, and comparing to a ground truth analytical expression; and 2) estimating MIND networks using a range of preprocessing configurations in N = 500 healthy UKB participants, namely smoothing, introduction of identical values due to quantisation of feature maps following storage in imprecise formats, analysis in native or common space and choice of vertex mesh resolution, parcellation choice, and feature selection, and benchmarking these networks by predicting age and sex, and correlating group mean networks with gene co-expression networks.

First, we used simulated Gaussian distributions with closed form expressions for KL divergence as a ground truth to evaluate the impact of five theoretical sources of bias in the KL divergence estimator:

1. the spatial **smoothness** or auto-correlation of input MRI feature maps;
2. **identical values**in input feature distributions, an under-appreciated consequence of certain MRI preprocessing operations;
3. vertex mesh resolution;
4. **region size differences** due to the choice of parcellation; and
5. the **covariance** between MRI features selected for multivariate MIND analysis.

Second, we highlight where these sources of bias emerge in a typical MIND network estimation pipeline using a normative MRI dataset from the UK Biobank (UKB) cohort. In the UKB dataset, we tested the hypothesis that the MRI preprocessing choices which were known (theoretically and by the prior analysis of simulated data) to incur least bias in the KL divergence estimator would also generate the most biologically valid estimates of brain morphometric similarity. Individual MIND networks estimated using various preprocessing pipelines were used to predict age and sex, and the sample mean MIND networks estimated using each preprocessing configuration were correlated with gene co-expression networks derived from the Allen Human Brain Atlas^31^. We largely confirmed our hypotheses that technically optimised MIND network phenotypes were also optimal in terms of the strength of their triangulation with biological and molecular brain phenotypes. Furthermore, we used the developing Human Connectome Project dataset^32^ to demonstrate that these bias sources can arise in a dataset collected with different MRI acquisition parameters and in a different developmental period. Finally, based on these analyses, we conclude with a set of recommendations to investigators wishing to use MIND in their own research.

## 2 Methods

### 2.1 Kullback-Leibler divergence and MIND

MIND is derived from the Kullback-Leibler divergence, *D_KL_*, an information theoretic measure for comparing two probability distributions or density functions, *P* and Q^33^:

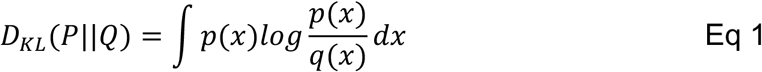

where *p*(*x*) and *q*(*x*) represent the probability densities in each distribution for a particular value *x* of the variable *X*. If *x* values have very different probabilities under distributions *P* and Q across a wide range of possible values of *X*, then the log ratios of probabilities will be large, and the KL divergence, calculated in aggregate as an integral or sum over all supported values of *X*, will be large.

The KL divergence can equivalently be expressed in terms of entropies:

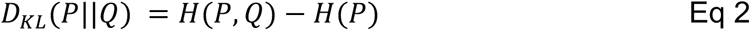

where *H*(*P*) is the entropy of *P* and *H*(*P*, Q) is the cross-entropy of *P* and Q. Eq 2 formalises the intuition that *D_KL_*(*P*||Q) is the additional information required to describe *P* when it is encoded using Q, that is, the information about *P* lost by predicting it from Q. Because the entropy of a distribution is always less than or equal to its cross-entropy with another distribution (Gibbs’ inequality^34^), KL divergence is bounded below by 0 and unbounded above, i.e., *D_KL_* ≥ 0.

Furthermore, KL divergence is asymmetric, i.e., *D_KL_* (Q||*P*) ≠ *D_KL_*(*P*||Q).

In many applications, KL divergence is symmetrised by Jeffreys’ transform to obtain

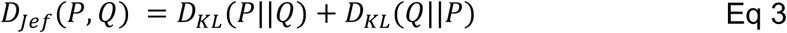

where *D_Jef_* is symmetric, non-negative, and unbounded above.

Symmetric Jeffreys’ divergence is inverted and normalised to convert it to a measure of similarity, MIND:

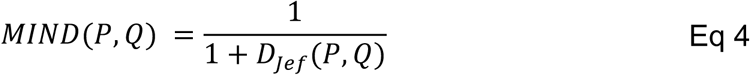

MIND is bounded between 0 and 1, where a high value indicates that a pair of regions have highly similar or undifferentiated morphometric feature distributions (low KL divergence) and a low value indicates that a pair of regions have highly dissimilar or differentiated distributions (high KL divergence).

Pairwise estimation of MIND between every possible pair of parcellated brain regions returns a symmetric similarity matrix, *M*, where each element or edge weight *M_i_*_j_ represents the morphometric similarity between regions *i* and >; and the sum of edge weights for each region ∑^j^ *M_i_*_j_ returns the weighted degree of the corresponding node in a network formulation of the MIND similarity matrix.

### 2.2 Estimators of Kullback-Leibler divergence

In practice, the continuous probability density functions underlying regional morphometric MRI data are not known *a priori* and must be estimated non-parametrically from a set of voxel or vertex values sampled in each brain region. Non-parametric KL divergence estimators for continuous distributions have been evaluated in the context of MIND networks^13^; the k-nearest neighbours (k-NN) estimator implemented in MIND^35^ consistently outperformed a histogram-based alternative when benchmarked against biological criteria.

The k-NN density estimator approximates the local probability density *p*(*x*) around a feature value *x* according to how close in value the k^th^ closest feature value is, which is the nearest value when k = 1^35^. Where samples are densely packed, this distance is small and the estimated density correspondingly high; where samples are sparse, the distance is large and the density low. A key advantage of this estimator is that it enables multiple features to be incorporated into one multivariate density estimation. Rather than computing Euclidean distance between neighbouring *values* of a single feature (e.g. cortical thickness) as a univariate measure of distance, the Euclidean distance between MRI feature *vectors* (e.g. comprising cortical thickness, sulcal depth, and mean curvature) is calculated.

The k-NN estimator can be used to estimate KL divergence directly by comparing the Euclidean distance *s*(*x*) between a feature vector *x* in *P* and its k^th^ nearest neighbour in Q to the Euclidean distance *r*(*x*) between *x* and its k^th^ nearest neighbour in *P*, excluding *x* itself. The probability densities local to *x*, for a feature vector of dimension *d*, are estimated as:

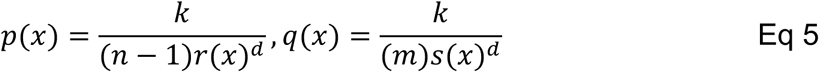

where *k* indicates which ranked neighbour is used to compute nearest neighbour distances and *n* and *m* denote the number of sample values of *X* used to represent the distributions *P* and Q respectively. Here, probability density is approximated by dividing the enclosed probability mass (*k*/(*n* − 1) for *P* and *k*/*m* for Q) by the volume of a *d*-dimensional ball. Since this volume is proportional to the radius raised to the power *d*, *r*(*x*)*^d^* and *s*(*x*)*^d^* stand in for enclosed volumes in *P* and Q, respectively. In the context of MIND network analysis, *n* and *m* relate to the number of voxels or vertices at which a multivariate MRI feature vector has been measured in regions *P* and Q, respectively.

Substituting these estimates of *p*(*x*) and *q*(*x*) into Eq 1 gives an expression for the k-NN estimator of the KL divergence:

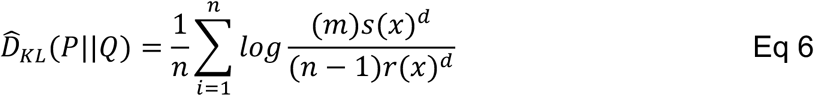

Eq 6 can be rewritten using mathematical log laws as:

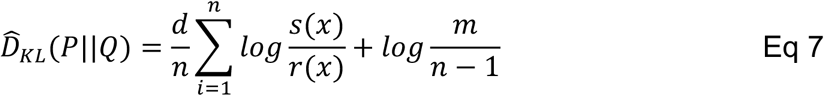

*D_KL_* can then be symmetrised by Jeffreys’ transform (Eq 3) and inverted (Eq 4) to generate a k-NN estimate of MIND similarity between the pair of brain areas represented by the multivariate feature distributions *P* and Q.

### 2.3 Factors influencing bias in the k-NN estimator

Bias refers to the systematic deviation between estimates and a known ground truth. Bias in the k-NN estimator has been explored in the context of KL divergence estimation^35^ as well as other information theoretic measures that use k-NN estimation as an intermediate step, such as differential entropy^36^ and mutual information^37^. Key sources of bias in the k-NN estimator that are of relevance to MRI data analysis are: (i) local uniformity of the *d*-dimensional sphere used to estimate local probability density; (ii) the number of statistically independent samples; and (iii) identical sample values.

#### 2.3.1 Local probability density uniformity

As noted, the k-NN estimator approximates probability densities around samples by dividing probability mass by the volume of the *d*-dimensional ball enclosing the k nearest neighbours. Critically, this approximation assumes that local probability densities around samples are uniform or approximately constant: non-uniformities can introduce bias. Non-uniformities can arise due to (i) high covariance between two or more dimensions, or (ii) differences in the marginal variances of one or more dimensions, both of which distort local neighbourhoods such that they take on elliptical rather than spherical shapes^38^. For instance, k-NN estimated mutual information between two Gaussian distributions is increasingly biased as the correlation between those variables grows: the joint density is progressively elongated along one axis, so the spherical neighbourhoods used by the estimator sample an increasingly anisotropic local density^37,39^. k-NN estimated KL divergence is also biased by non-uniformities in probability density. KL divergence between two Gaussian distributions is biased at low sample sizes when there are differences in distributional support overlap between distributions, i.e., when there is a zone of probability density in one distribution with low or absent probability density in the other distribution^35^. For a sample from *P* located in such a zone, the ball must inflate considerably before reaching its k^th^ nearest neighbour in Q, and the density it encloses is correspondingly non-uniform, biasing KL most severely at low sample sizes^35^.

#### 2.3.2 Sample size

k-NN density estimates converge to the ground truth value, i.e. bias decreases, as the number of statistically independent samples used by the estimator is increased^35,37,40,41^. Therefore, for small finite sample sizes, k-NN estimates are biased. In neuroimaging data, vertex and voxel-level data are spatially autocorrelated^42^, so the nominal sample size, i.e. the total number of vertices or voxels, overstates the number of statistically independent samples available to the k-NN KL estimator^43^.

Critically, the sample size required for convergence of the k-NN KL estimator is dependent on local non-uniformities, described in **Methods 2.3.1**. Where the local uniformity assumption is more strongly violated, a convergence of k-NN estimates with the ground truth value requires more samples. For example, when the support overlap between two distributions is low, i.e. a substantial part of the mass in *p*(*x*) lies where *q*(*x*) is barely sampled, *D_KL_* (*P*||Q) requires far more samples to converge than the reverse direction *D_KL_* (Q||*P*)35. Bias therefore grows as the available sample size falls relative to the degree of local uniformity. This concept has been applied to importance sampling, i.e. using samples from one distribution to estimate expectations under another: the sample size required for accurate estimation grows with the true divergence between the two distributions^44^.

#### 2.3.3 Identical sample values

The k-NN estimator requires sample values to be non-identical because identical values concentrate mass at a single point, resulting in nearest neighbour distance, *r*(*x*) or *s*(*x*), equalling 0 and *log s*(*x*)⁄*r*(*x*) in Eq 7 being undefined. This is problematic for biological data, in which identical values may exist due to limitations on measurement precision or data preprocessing operations^45^. To navigate this issue, the current implementation of the k-NN KL estimator in MIND ignores samples whose nearest neighbour distance in either distribution is 0.

### 2.4 Evaluation of KL and MIND bias using simulated Gaussians

In light of the theoretical causes of bias in k-NN-based estimators, we hypothesised that operations or choices made during typical neuroimaging pipelines could lead to introduction of bias during KL and MIND estimation. To investigate this, we simulated the effects of various MRI preprocessing operations on bias in KL and MIND. Bias in KL and MIND were computed by comparing ground truth values calculated from a closed form expression for KL divergence for Gaussian distributions with the equivalent values estimated using the k-NN method.

More precisely, for two Gaussian distributions, *P* and Q, parametrised by known mean and variance, such that *P*∼*N*(*μ_P_*, *σ_P_*) and Q∼*N*(*μ*_Q_, *σ*_Q_), a closed form expression for KL divergence between them can be expressed as a function of their means and variances:

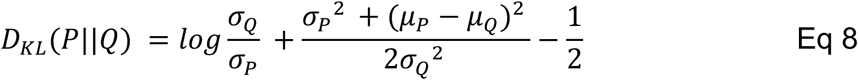

Eq 8 symmetrises to give:

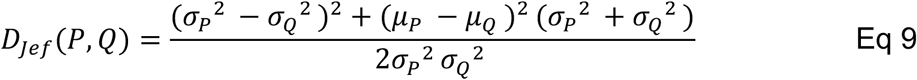

The expression in Eq 9 makes explicit the contributions of both variance mismatch and mean separation towards symmetric KL divergence. The closed-form solution to KL can also be used to compute ground-truth KL divergence between multivariate Gaussian distributions^46^.

Points were sampled from distributions *P* and Q and the symmetric form of k-NN estimated KL divergence *D_Jef_*(*P*, Q) was calculated. The percentage bias between estimated and ground truth KL divergence, *D_Jef_*(*P*, Q), was defined as:

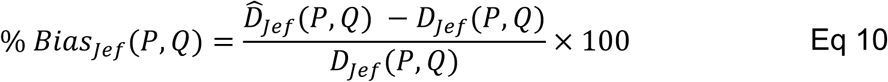

Likewise, the bias in estimation of MIND was calculated as:

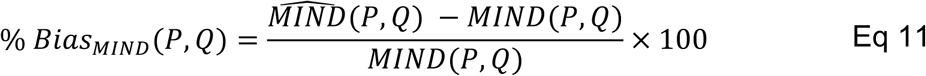

where MIND is the estimated value, inverted from k-NN estimated KL, and *MIND* is the true value, inverted from the closed form expression of KL (Eq 9).

Computing bias while modulating the parameters of input feature distributions permits a systematic investigation of bias in KL and MIND estimation. We used this bias assessment framework to explore the effects of five vertex map properties on absolute values or bias in KL and MIND: (i) smoothness; (ii) identical values introduced by quantisation following storage in imprecise formats; (iii) resolution; (iv) parcellation region size; and (iv) feature covariance (**Table 1**). In each case, samples from randomly initialised Gaussian distributions were modified to express each vertex map property to varying extents, and absolute values or bias in KL and MIND were computed for each extent of modification. This procedure was repeated and averaged over 50 pairs of Gaussian distributions with means and variances randomly sampled from *μ* ∈ [1.7, 2.9], *σ* ∈ [0.4, 1.4], reflecting empirical ranges of regional means and variances from the cortical thickness map of a representative adult brain. Simulations were designed to recapitulate effects of preprocessing steps on vertex maps, rather than to specifically isolate bias sources enumerated in **Methods 2.3**. In some cases, simulations influence multiple biasing factors at once (see **Results**).

**Table 1:** Simulation experiments and variable parameters. Simulated properties are presented in the order in which they may be implicitly introduced or explicitly applied during typical MRI preprocessing pipelines (**Figure 1**). The number of feature dimensions reflects the dimensionality of the simulated data (N = 1: univariate, N > 1: multivariate), and the number of spatial dimensions indicates the number of spatial dimensions onto which points were sampled; for example, when N spatial dims = 2, points were sampled onto a 2-dimensional grid. In each experiment, 30 evenly spaced values were selected from the variable parameter ranges specified, giving 900 combinations. In the smoothing, quantisation, vertex mesh resolution and feature covariance experiments, points were drawn from a pair of ground truth Gaussian distributions with known parameters, using 4000 samples per distribution. Either the sampled points were modulated by varying degrees of smoothing, quantisation, or resampling, or the covariance between features within the ground truth distributions was itself modulated. In these cases, the generating, ground-truth, Gaussian distributions are specified, so KL and MIND estimated from modulated samples can be expressed as bias relative to analytic ground truth KL and MIND. In the parcellation experiment, a spatially autocorrelated Gaussian random field was generated across a 2-dimensional grid, and non-overlapping patches were grown from random seed points to cover a specified percentage of the grid. Because points are not drawn from ground truth Gaussian distributions, but are instead defined by patch size, absolute KL and MIND were quantified rather than bias relative to an analytic ground truth. In each experiment, performance was averaged over 50 unique Gaussian or Gaussian random field specifications.

| Simulated property | Outcome | N feature dims | N of spatial dims | Variable parameter | Variable parameter ranges |
| --- | --- | --- | --- | --- | --- |
| Smoothness | KL and MIND bias | 1 | 2 | Smoothing of points sampled from ground truth distributions | $FWHM \in [0,5]$ |
| Identical values due to quantisation | KL and MIND bias | 1 | 0 | Quantisation of points sampled from ground truth distributions | $Q \in [10^{-4}, 10^{-2}]$ |
| Vertex mesh resolution | KL and MIND bias | 1 | 1 | Extent of resampling of points sampled from ground truth distributions | $E \in [-0.8, +0.8]$ |
| Parcellation | Absolute KL and MIND | 1 | 2 | Percentage of grid coverage | $\% \in [2,50]$ |
| Feature covariance | KL and MIND bias | 5 | 0 | Covariance of ground truth multivariate distributions | $\Sigma \in [0,0.9]$ |

### 2.5 Triangulation of MIND networks against biological and molecular phenotypes

We hypothesised that vertex map properties that generated high bias in KL and MIND in simulated data would reduce the biological validity of MIND networks when applied to input feature maps. To test this, we validated MIND networks generated using a range of preprocessing parameters across three analyses: age and sex prediction, and correspondence with gene co-expression similarity networks.

#### 2.5.1 Age and sex prediction

Prediction of age and sex was chosen as validation criteria on the basis that inter-areal similarity in cortical structural properties, measured using MIND, is strongly influenced by biological differences due to age and sex, as demonstrated in previous work^4,11–13^. Under this framework, MIND network specifications that more accurately predict age and sex may be more biologically valid constructs^47^.

We trained ridge regression models to predict age and ridge classifier models to predict sex using all edge weights from MIND networks. Ridge models were selected as they are computationally tractable to run on many MIND network configurations. Models were trained and evaluated on 50 random splits of the data, with each training fold comprising 75% of the data (N = 375). To attenuate confounding, edge weights were first residualised by regression on estimated total intracranial volume, Euler index, and a measure of head motion derived from T1-weighted images (**Methods 2.6.1**)^48^. Within each fold, the train and test sets were residualised using a multiple linear regression trained on the training set only, and residualised edge weights were standardised using a z-scorer fitted on the training set only, to avoid data leakage^49^. Within each training fold, leave-one-out cross-validation was also used to select optimal regularisation values from 50 logarithmically spaced values in the interval [10^−3^, 10^6^]. The optimal ridge model was trained on the full training set and evaluated on the test set using either Pearson correlation and mean absolute error (MAE), for regression models, or the area under the receiver operating characteristic curve (AUROC), for classifier models.

To assess whether there were significant differences in fold-wise prediction accuracies between MIND network preprocessing configurations, we used the non-parametric Friedman test, treating cross-validation folds as paired observations given that identical training/test splits were used across all preprocessing configurations. To assess for significant pairwise differences between conditions, we used the Nemenyi post-hoc test which inherently controls the family-wise error rate.

#### 2.5.2 Gene co-expression correspondence

Several studies have demonstrated high correlations between MRI structural similarity networks and gene co-expression networks^4,7,13^, with inter-regional similarity at the macroscale thought to be reflective of, or emerging because of, inter-regional similarity at the transcriptomic or genetic level^2^. This expectation reflects Cheverud’s conjecture^50^, whereby phenotypic covariance between areas, whether in terms of gene co-expression or MIND similarity, should be driven by genetic correlation between areas, and therefore gene co-expression and MIND similarity should be correlated with each other.

We computed the Pearson correlation value between edges of group mean MIND networks and equivalently parcellated gene co-expression networks. Because the AHBA comprises only six donors, we assessed the sensitivity of these correlations to donor sampling by recomputing the gene co-expression network for all 63 non-empty combinations of donor subsets, and summarised correlation values as the mean ±1 standard deviation for each donor subset size.

### 2.6 MRI and gene expression data preprocessing

#### 2.6.1 UK Biobank (UKB)

We used T1-weighted and diffusion MRI data from the UKB, a large-scale association study of predominantly healthy adults in the UK. Acquisition, volumetric preprocessing, and quality control of MRI data has been described previously^51,52^. The acquisition parameters used were:

- T1-weighted (T1w): 3D magnetisation-prepared rapid gradient echo sequence, 1.0×1.0×1.0 mm, 208×256×256, R=2, TI/TR=880/2000 ms.
- T2-weighted fluid attenuated inversion recovery (T2 FLAIR): 3D sampling perfection with application-optimized contrasts using different flip angle evolution (SPACE) sequence, 1.0×51.0×1.0 mm, 192×256×256, R=2, PF 7/8, fat saturation, TI/TR = 1800/5000 ms.
- Diffusion MRI (dMRI): multiband echo planar imaging sequence, 2.0×2.0×2.0 mm, 104×104×72, MB=3, R=1, fat saturation, b=0(5x + 3x phase-encoding-reversed), 1000(50x), 2000(50x).

We selected a representative subset of 500 healthy individuals for analysis (mean age ± 1 standard deviation = 61.2 ± 7.2 years, N female = 250). Individuals were deemed healthy if they had no self-reported cancers or non-cancer illnesses. Volumetric image preprocessing, surface interpolation, and parcellation were performed using FreeSurfer 6.0.0^53^. T2w images were used, where available, to enhance the accuracy of pial surface reconstruction. Five macrostructural features were derived from T1w scans as part of the standard FreeSurfer pipeline: cortical thickness (CT), surface area (SA), volume (vol), sulcal depth (SD) and mean curvature (MC). Preprocessing of dMRI data has been described previously^52^: first, diffusion data were corrected for eddy currents, head motion, and outlier slices, after which diffusion tensors were fit, generating fractional anisotropy (FA) and mean diffusivity (MD) images; subsequently, Neurite Orientation Dispersion and Density Imaging (NODDI) modelling was performed, generating orientation dispersion (OD), intracellular volume fraction (ICVF), and isotropic volume fraction (ISOVF) images. Additional preprocessing has been described previously^54^: dMRI images were co-registered to T1w space using the `bbregister` command in FreeSurfer, and surface interpolation was subsequently performed using the `volume-to-surface-mapping` command in Connectome Workbench^55^, resulting in dMRI feature maps evaluated in native surface space.

Surface features evaluated in native subject space were subsequently projected to a common template, fsaverage6 (40,962 vertices per hemisphere), using FreeSurfer’s `mri_surf2surf`^56^. Feature maps in fsaverage6 space were aligned to three other common space templates: fsaverage, fsaverage5, and fsaverage4 (163,842, 10,242, and 2,562 vertices per hemisphere, respectively), using the neuromaps `transforms.fsaverage_to_fsaverage` function with barycentric interpolation^57^. We avoided directly upsampling from native space to the higher resolution fsaverage template using FreeSurfer’s `mri_surf2surf`, as this function only supports nearest neighbours mapping, which introduces identical values when upsampling^56^. This procedure resulted in 5 macro- and 5 microstructural surface-based morphological maps evaluated in native space and three common spaces.

Feature maps in native and common space were parcellated using two hierarchically nested cortical parcellation schemes based on the Desikan-Killiany (DK)^58^ and Human Connectome Project Multi-Modal (HCP)^59^ parcellations. DK68 and DK318 comprise 34 and 159 symmetric cortical areas per hemisphere, respectively, with the fine-grained DK318 parcellation generated by hierarchically subdividing the coarse-grained DK68 template into approximately equally sized cortical areas^60^. The fine-grained HCP360 template, comprising 180 cortical areas per hemisphere^59^, was coarse-grained to 23 bilateral cortical areas to form the HCP46 parcellation^18^. After filtering for invalid feature values, 24.8% of subjects had zero vertices remaining in at least one of their hippocampal regions when parcellated using HCP360. We therefore excluded these regions from both hemispheres of the HCP360 parcellation, leaving 179 regions per hemisphere. Finally, we used surface-based alignment to project these atlases to each subject’s native space.

Analyses of brain structure and function can be confounded by scan quality, head motion, and brain size^48^. To directly control for these confounds in downstream analyses, we used a number of QC metrics provided by UKB: Euler number, defined as the total number of holes in left and right hemispheric cortical surface reconstructions prior to topological correction^61^ as a metric of T1w and T2w scan quality; a proxy of head motion derived from linear regression-based prediction of manually rated T1w scan motion artefacts^48^; and estimated total intracranial volume (eTIV).

#### 2.6.2 Developing Human Connectome Project (dHCP)

We used neonatal brain MRI data from the dHCP^32^ to illustrate the prevalence of bias sources in a dataset differing in acquisition and lifespan stage to the UKB. T2-weighted MRI scans were acquired shortly following birth for N = 752 neonates, aged 26-43 weeks post-menstrual age (**Table 2**) using a multi-slice fast spin-echo sequence: TR=12s, TE=156ms, SENSE factor = axial 2.11, sagittal 2.66; in-plane resolution 0.8×0.8 mm^2^; 1.6 mm slices. These data were processed using a pipeline of cortical surface reconstruction and measurement of macrostructural MRI features (cortical thickness, mean curvature, sulcal depth, surface area, and volume)^62^, then regionally parcellated using the Melbourne Children’s Regional Infant Brain Surface (MCRIB-S) atlas^63^.

**Table 2:** Developing Human Connectome Project MRI cohort demographic and clinical data. PMA: post-menstrual age. std: standard deviation.

| Characteristic | N (%) or mean ± std |
| --- | --- |
| Total subjects | 752 |
| Sex (M/F) | 407 (54%) / 345 (46%) |
| Twin subjects | 100 (13%) |
| PMA at birth (weeks) | 37.8 ± 4.13 |
| PMA at scan (weeks) | 39.8 ± 3.53 |
| Birth weight (kg) | 1.68 ± 0.92 |
| Radiology score (/5) | 1.91 ± 1.19 |

#### 2.6.3 Allen Human Brain Atlas of cortical gene expression

We used spatially resolved bulk transcriptomic data from the Allen Human Brain Atlas (AHBA^31^), which sampled the brains of six post-mortem human donors (mean age = 42 years, N female = 2). AHBA data from the left cortical hemisphere was preprocessed using Abagen version 0.1.3^64^, after which a validated preprocessing pipeline^65^ was applied, comprising: intensity-based filtering using a threshold of 0.5; selection of probes with high differential stability across donors; mapping of samples to the nearest parcellated region; and normalisation of gene expression within samples and within donors using the scaled robust sigmoid function. Transcriptional similarity between each pair of regions was computed using an angular similarity metric suitable for computing similarities between highly correlated, high-dimensional gene expression vectors^4,66^. This procedure was repeated using the same 4 parcellations as were used for MRI data (**Methods 2.5.1**) and all 63 non-empty donor subsets, to assess stability of correlations with MIND networks (**Methods 2.5.2**).

### 2.7 Statistical approaches

#### 2.7.1 Resampling to remove identical values

We sought to remediate the effects of identical values via a resampling procedure. This procedure involved estimating a probability density function (PDF) from the identical value-containing distribution using kernel density estimation (KDE), performed using SciPy’s `stats.gaussian_kde()` function with default parameters^67^. Independent and identically distributed (i.i.d.) samples were then drawn from this PDF to generate a distribution of samples with statistics approximately matching the original distribution, though without identical values.

We assessed how efficacious this resampling procedure was at recovering original MIND network edge structure in simulated data. For 10 Gaussian distributions with parameters randomly chosen from *μ* ∈ [1.7, 2.9], *σ* ∈ [0.4, 1.4], the same empirically representative range of values used in other simulation experiments, MIND was computed before (Original), after quantisation to an extent randomly chosen from the range Q ∈ [10^−4^, 10^−2^] (Quantised), and after resampling the identical value-containing distributions (Resampled). We assessed the edge-wise correspondence between the Original and Resampled networks using Spearman correlation and mean absolute error (MAE) over a range of resampled sample sizes.

#### 2.7.2 Resampling extent

To quantify the extent to which simulated data or vertex maps had been up- or downsampled in experiments relating to vertex mesh resolution, we computed resampling extent as the proportional difference in sample number:

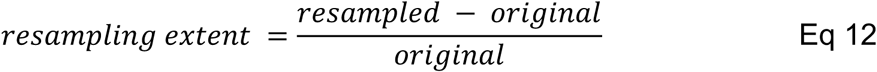

where ‘original’ refers to the number of samples in simulated or empirical data before resampling and ‘resampled’ refers to the target number of samples in the resampled data after resampling. A resampling extent of −0.5 indicates a 50% decrease in the number of samples after resampling, +0.5 indicates a 50% increase, and 0 indicates no change.

#### 2.7.3 Gini coefficient

To quantify region size homogeneity across parcellations, we used the Gini coefficient^68^, which varies from 0 (perfectly homogeneous, i.e. all regions contain the same number of vertices) to 1 (perfectly inhomogeneous, i.e. one region contains all vertices and the remainder contain none). The Gini coefficient is calculated as the mean pairwise difference in number of vertices across all cortical regions, normalised by the mean number of vertices across regions:

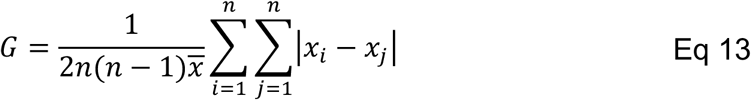

where n is the number of regions, *x_i_* and *x*_j_ are the number of vertices in region *i* and *j*>, respectively, and *x* is the mean number of vertices in a region.

#### 2.7.4 Principal component analysis

Based on theoretical understanding and simulation results that KL divergence is impacted by feature covariance, we orthogonalised MRI features using principal component analysis (PCA) of vertex maps. We computed principal components (PCs) using two distinct feature sets: 5 macrostructural features or all 10 macro- and microstructural features.

To ensure that components were identically ordered and oriented across subjects, a single standardisation and PCA model were fit across the full sample so that all subjects shared a common basis. Because concatenating each individual’s vertex-by-feature matrix would be prohibitively memory intensive, both models were fit incrementally in batches of subjects on vertex-wise data evaluated in fsaverage5 space: a z-scorer was first fit across batches with scikit-learn’s `StandardScaler.partial_fit()`; each subject’s matrix was then standardised using the fitted z-scorer; and finally a PCA model was fit across batches using `IncrementalPCA.partial_fit()`^69^. Individual raw vertex-by-feature dataframes, evaluated in native space, were subsequently z-scored and projected to component space using the scaling parameters and component loadings estimated on the full sample.

To assess the extent to which the loadings of morphometric features on principal components aligned when computed across different population subsamples or cohorts, we computed Tucker’s congruence coefficient, equivalent to the cosine similarity following L2-normalisation^70^, between each pair of components, and selected component pairings which maximised the summed absolute congruence using the Hungarian algorithm^71^. Absolute congruence, |*Φ*|, was retained for each pair, with |*Φ*| ≥ 0.95 indicating that components could be considered equivalent^70^.

## 3 Results

### 3.1 Image smoothness

#### 3.1.1 Sources of smoothness

Diverse cortical properties naturally demonstrate high spatial smoothness, or autocorrelation, whereby spatially proximal cortical areas show more similar values than spatially distant areas^42^. However, routine steps in structural MRI processing pipelines can additionally increase the extent of smoothness of MRI-derived morphometric features: MRI images can be deliberately smoothed during preprocessing to enhance signal-to-noise^72,73^, or non-deliberately smoothed during surface reconstruction, interpolation between cortical surfaces, or projection from volumetric to surface space.

#### 3.1.2 Effects of smoothness on KL divergence and MIND in simulated Gaussian distributions

We hypothesised that smoothing would increase k-NN estimator bias via two channels. First, smoothing increases the dependence between neighbouring vertices, reducing the number of statistically independent samples for KL estimation (**Methods 2.3.2**). Second, smoothing decreases within-region variance, reducing the degree of distributional support overlap on average, and potentially violating the local uniformity assumption of the k-NN estimator (**Methods 2.3.1**).

To investigate the effects of smoothness on bias in KL divergence and MIND, we simulated smoothing by sampling from Gaussian distributions onto 2-dimensional grids, smoothing the grids using Gaussian kernels with independently varied full width at half maximum (FWHM), and computing bias in estimated KL and MIND relative to a ground truth, closed-form solution. Simulations revealed that greater smoothing resulted in positively biased KL divergence values and negatively biased MIND values, with a maximum absolute MIND bias of −73% observed when both grids were maximally smoothed (**Figure 2a-b**).

**Figure 2:**
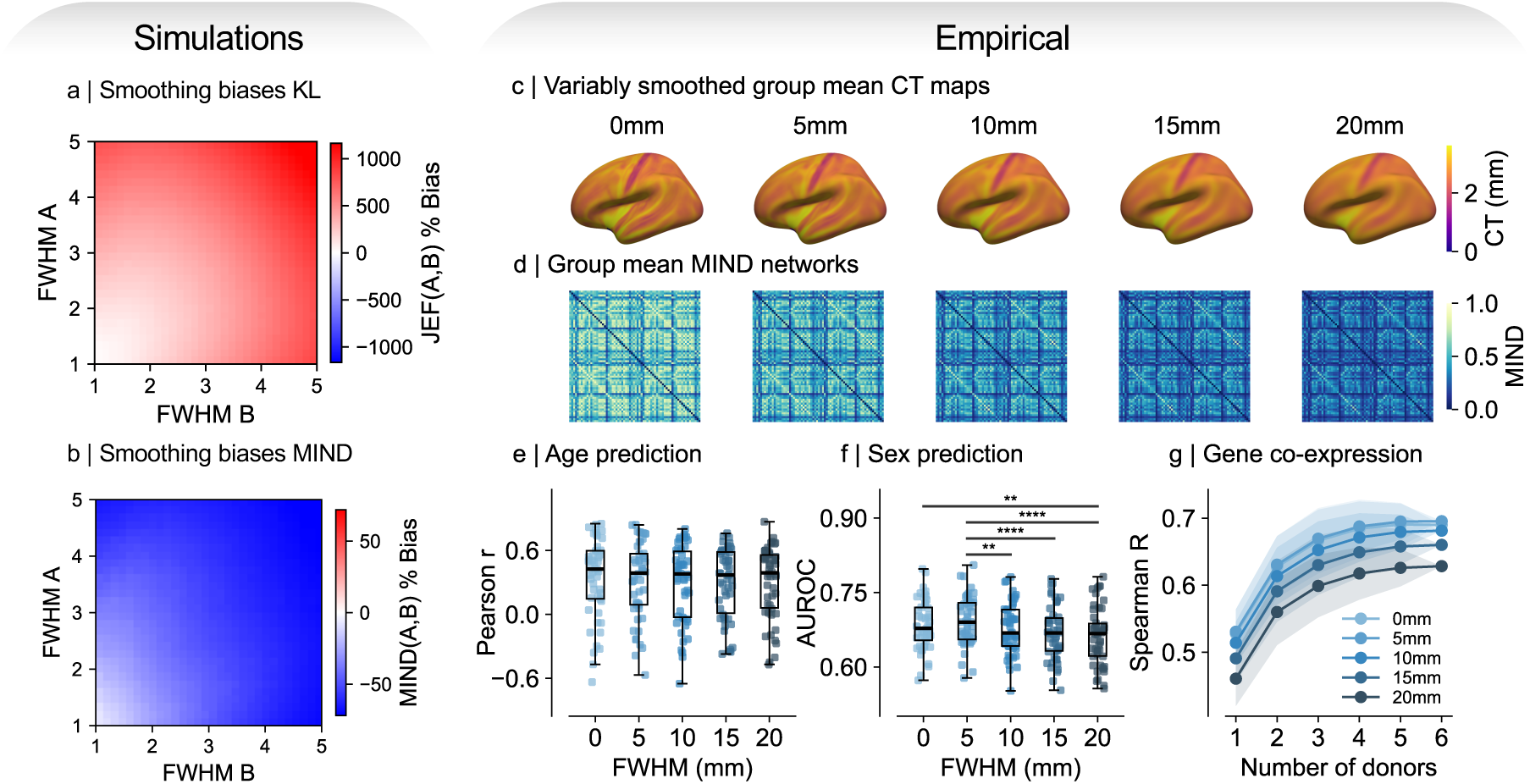
Smoothing biases KL divergence estimation and reduces the biological validity of MIND. Percentage bias in **a** symmetric KL divergence (JEF) and **b** MIND as smoothness of each distribution was independently varied. **c** Group mean cortical thickness map smoothed using a Gaussian kernel with FWHM = [0, 5, 10, 15, 20] mm. **d** Group mean MIND networks for each smoothing condition. **e** Increasing smoothing did not significantly impact the accuracy of age predictions. **f** Increasing smoothing significantly decreased the accuracy of sex classification. In **e-f**, bars indicate pairwise significant relationships following the Nemenyi post-hoc test, with * p < 0.05, ** p < 0.01, *** p < 0.001, **** p < 0.0001. **g** Correlations between group mean MIND networks and gene co-expression networks decreased as input cortical thickness maps were increasingly smoothed. Lines show the mean and shaded areas show ±1 standard deviation of correlation values across all non-empty combinations of donors used to estimate gene co-expression networks.

#### 3.1.3 Effects of smoothness on MIND in structural MRI data

To evaluate the effects of vertex map smoothness on MIND networks estimated from empirical data, we smoothed native space cortical thickness maps using a 2D Gaussian kernel with FWHM = [0, 5, 10, 15, 20] and estimated univariate MIND networks from these maps. MIND networks estimated at these five smoothness levels were validated using age and sex prediction and correlations with gene co-expression networks.

Greater smoothing of cortical thickness maps (**Figure 2c**, **Supplementary Figure 1a**) reduced intra-areal variances of cortical thickness (**Supplementary Figure 1b**), increased the KL divergence between areal distributions of cortical thickness (**Supplementary Figure 1c**), globally decreased MIND edge weights (**Figure 2d**, **Supplementary Figure 1d**), and increased the variability of edge weights across subjects (**Supplementary Figure 1e**). As smoothing of cortical thickness maps was increased, there were no significant differences in age prediction accuracies (omnibus

Friedman p = 0.08, **Figure 2e**), while sex classification accuracies significantly decreased (omnibus Friedman p < 0.001, **Figure 2f**). The magnitude of correlations between MIND networks and gene co-expression networks decreased as cortical thickness maps were increasingly smoothed (**Figure 2g**).

#### 3.1.4 Summary of bias risks and mitigations for smoothing

In summary, increases in estimator bias due to smoothing, that were predicted *a priori* and demonstrated in simulated data, may also underpin the reduced biological validity of resulting MIND networks. We therefore recommend against application of additional smoothing to vertex maps to enhance signal-to-noise prior to MIND network estimation.

### 3.2 Identical values

#### 3.2.1 Sources of identical values

Identical values can arise in cortical surface feature maps measured with finite precision as part of normal biology, for example in the smooth cortex of early neonatal brains or lissencephalic non-human species. We analysed the prevalence of identical values in the dHCP dataset and identified age- and feature-specific trends (**Supplementary Figure 3a**). Younger neonates had a higher proportion of identical values in cortical thickness, volume, and, to a lesser extent, surface area, which decreased as neonates matured, reflecting the normal developmental transition from a highly homogeneous, smooth cortical surface to an increasingly complex and gyrified surface.

Identical values are also inadvertently introduced during typical image preprocessing pipelines. For example, identical values can be introduced when storing data in efficient but imprecise storage formats, such as INT16 or INT8, which can hold 2^15^ = 32,768 or 2^7^ = 128 unique values per sign, respectively. Conversion of relatively precise data to imprecise formats quantises the data such that unique values take one of a smaller number of discrete values. To illustrate this in a typical neuroimaging pipeline, volumetric T1w images are converted to UCHAR format, storing 256 unsigned integers, by the `mri_convert --conform` command as part of FreeSurfer’s `recon-all` pipeline^74^. Identical values can also be introduced when upsampling a voxel image or vertex map from a low to higher resolution using nearest neighbours interpolation, as may occur when analysing subjects on a high-resolution template, such as fsaverage. Nearest neighbours interpolation assigns multiple target voxels or vertices the value of their nearest source voxel or vertex, thereby introducing identical values. This is the default behaviour of FreeSurfer’s `mri_surf2surf` function^56^.

#### 3.2.2 Effects of identical values on KL divergence and MIND in simulated Gaussian distributions

Since the k-NN estimator relies on comparing distances between points and their k^th^ nearest neighbours, identical feature values within or across distributions pose a mathematical invalidity: the log ratio between the nearest neighbour distances will result in either a *log*(0) or *log*(1/0) (**Methods 2.3.3**). This is most problematic for univariate MIND networks, since the chance of two samples sharing exactly identical values drops sharply as more features are added. To enable probability estimation, the current implementation of MIND ignores these samples, setting its contribution to the log-ratio of distances to 0. This approach does not entail significant alteration of input distributions when the proportion of identical sample values they contain is low, however, for distributions containing a high proportion of identical values, systematic sample exclusion can produce drastic changes. Since identical feature values provide key information about highly probable values within each distribution, we hypothesised that removing them in bulk may remove key signal from the k-NN probability estimation, introducing bias.

To assess the impact of identical feature values on bias in KL divergence and MIND estimation, we simulated identical values by drawing samples from Gaussian distributions with independently varied degrees of quantisation, excluded identical samples in line with the standard MIND procedure, and computed bias in estimated KL and MIND relative to a ground truth, closed-form solution. These simulations revealed that greater quantisation, synonymous with higher proportions of identical values, led to positive bias in estimated KL and negative bias in MIND, with a maximum absolute MIND bias of −0.31% observed when both distributions were maximally quantised (**Figure 3a-b**). Further simulations revealed that the influence of input data quantisation on MIND networks was strongly attenuated by our resampling procedure, and using N ≈ 4000 resampled points yielded corrected MIND networks that were highly correlated (Pearson r = 0.98) with original MIND networks prior to artificial introduction of identical values by quantisation (**Supplementary Figure 2**). All downstream analyses estimating MIND from resampled data therefore use N = 4000 resampled points.

**Figure 3:**
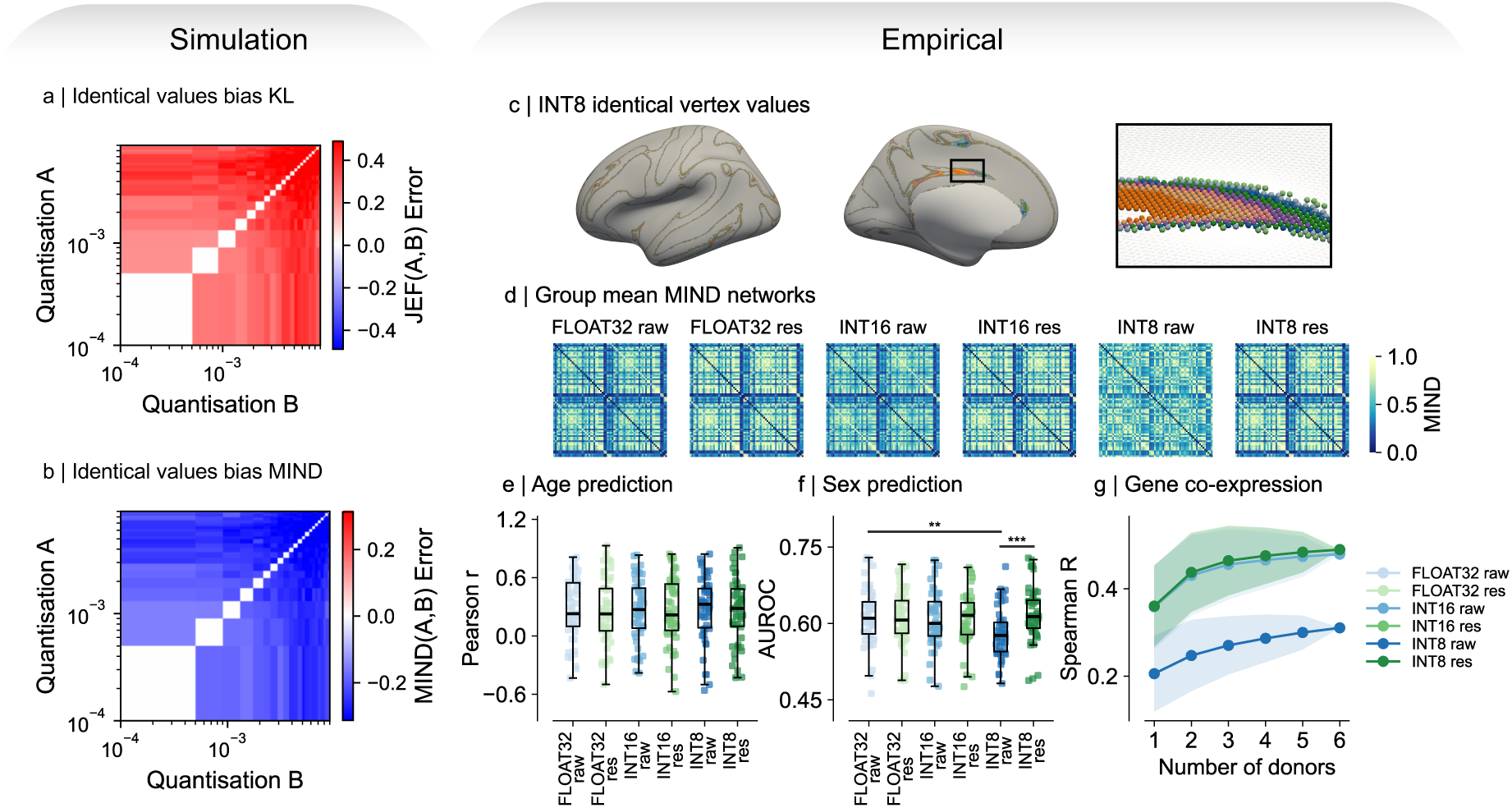
Identical values bias KL divergence estimation and reduce the biological validity of MIND. Percentage bias in **a** symmetric KL divergence (JEF) and **b** MIND as the degree of quantisation of each distribution was independently varied. **c** Group mean SD map quantised to reflect storage in INT8 format, with the spatial locations of N = 10 identical vertex values highlighted. **d** Group mean MIND networks generated from SD maps that had been quantised to varying extents (raw) and optionally resampled (resampled). **e** Quantisation did not significantly impact the accuracy of age predictions. **f** High quantisation (INT8) significantly decreased the accuracy of sex classification (FLOAT32 raw vs. INT8 raw: p = 0.001), which was recovered by resampling (INT8 raw vs. INT8 resampled: p < 0.001). In **e-f**, bars indicate pairwise significant relationships following the Nemenyi post-hoc test, with **\*** p < 0.05, ** p < 0.01, *** p < 0.001, **** p < 0.0001. **g** Correlations between group mean MIND networks and gene co-expression networks were reduced for MIND networks generated from INT8 quantised SD maps and recovered by resampling. Lines show the mean and shaded areas show ±1 standard deviation of correlation values across all non-empty combinations of donors used to estimate gene co-expression networks.

#### 3.2.3 Effects of identical values on MIND in structural MRI data

To explore the effects of quantisation-induced identical values on MIND network validity in empirical data, sulcal depth maps in FLOAT32 format were artificially quantised to reflect the precision of INT16 or INT8 data storage formats. Applying this quantisation procedure to the group mean SD map, we found that the percentage of values with at least one identical counterpart across the cortex was 8.7% in FLOAT32, 72.3% in INT16, and 99.8% in INT8 format, with larger regions containing a larger proportion of these values according to their coverage of the cortex (**Supplementary Figure 3b**). The spatial locations of 10 regularly repeating vertex values in the INT8 map are shown in **Figure 3c**.

Univariate MIND networks were computed from the non-quantised (FLOAT32) and variably quantised sulcal depth maps (INT16 and INT8). To assess the efficacy of resampling at mitigating bias due to identical values, three additional MIND networks were computed from these variably quantised sulcal depth maps after resampling with N = 4000 samples per cortical area, yielding six networks per subject (**Figure 3d**). Resampling was performed by calling `compute_MIND_networks()` with the `resample=True` flag. All resulting networks were highly correlated at the edge level (Pearson r ≥ 0.97), except for the non-resampled INT8 networks (Pearson r = 0.47-0.54, **Supplementary Figure 3c**). In these networks, regional weighted degree was tightly non-linearly related to the regional number of vertices, suggesting that MIND network structure was driven primarily by regional size (**Supplementary Figure 3d**).

MIND networks generated from INT8 sulcal depth maps produced age predictions that were not significantly different from other network specifications (omnibus Friedman p = 0.57, **Figure 3e**) but demonstrated significantly less accurate predictions of sex than networks generated from unquantised data (Nemenyi p = 0.001, **Figure 3f**). Group mean INT8 networks also had much weaker correlations with gene co-expression (**Figure 3g**). Relative to networks generated from quantised INT8 sulcal depth maps, resampling improved the accuracy of sex predictions (Nemenyi p < 0.001) and increased correlations with gene co-expression, highlighting the efficacy of this approach at mitigating bias due to identical values.

#### 3.2.4 Summary of bias risks and mitigations for identical values

In summary, identical values bias KL divergence in simulated data and can reduce the biological validity of resulting MIND networks in the univariate case. MRI data can naturally contain identical values, though we observed insufficient proportions (<15%) to cause appreciable bias in neonatal brain data (**Supplementary Figure 3a**). Researchers should be aware that common MRI preprocessing steps, such as storage in imprecise data storage formats and upsampling with nearest neighbours interpolation, can introduce identical vertex values. However, resampling can mitigate this bias, as we demonstrate using simulated and empirical data.

### 3.3 Analysis space and vertex mesh resolution

#### 3.3.1 Sources of variation in analysis space and vertex mesh resolution

A critical MRI preprocessing decision is whether to analyse data in native space, preserving subject-specific geometry, or in common space, enhancing inter-individual comparability by projecting subject-specific data onto a common template. Native-space analysis introduces inter-individual variability in vertex counts, with total vertex counts scaling with surface area. We found that this variability is pronounced in the developmental dHCP cohort, where vertex counts rise steeply with age alongside cortical expansion, but is comparatively stable in the adult UKB cohort (**Supplementary Figure 4**). Resampling to a common vertex mesh, e.g. fsaverage6, ensures that subjects have the same total vertex counts. If subjects have widely varying native space vertex counts, this may involve variably up- or downsampling subjects, as we observed in the dHCP cohort (**Supplementary Figure 4a**). The extent of resampling applied therefore varies systematically across subjects, which we show below is itself a source of bias in KL and MIND estimation.

#### 3.3.2 Effects of vertex mesh resolution and resampling on KL and MIND estimation in simulated Gaussians

Vertex mesh resolution can be increased or decreased by barycentric interpolation, in which each resampled value is a weighted average of the source vertices mapping onto it. We note that upsampling can be additionally performed by nearest-neighbour interpolation, though this introduces identical values which we have shown biases the k-NN estimator (**Figure 3a-b**), and downsampling can additionally be performed by dropping samples, though this can only serve to increase bias in the k-NN estimator, which converges to ground truth as the number of independent samples is increased (**Methods 2.3.2**). We therefore focus on barycentric interpolation, which is the default interpolation method used in several software packages, including in the surface-to-surface mapping tools provided by Connectome Workbench^55^.

Barycentric interpolation acts asymmetrically on the two nearest-neighbour distances entering Eq 7, and this asymmetry generates bias. Within-region distances, *r*(*x*), are sensitive to local spatial sampling because for morphological feature maps, which are highly spatially autocorrelated, the k^th^-nearest neighbour (i.e. the neighbouring feature value) of a vertex is typically also its spatial neighbour. Conversely, for between-region distances, *s*(*x*), the k^th^-nearest neighbour of a vertex is generally not its spatial neighbour and is therefore largely unaffected by changes in local sampling density.

Consequently, upsampling by interpolation is predicted to shrink *r*(*x*), because each interpolated vertex is a weighted average of its source vertices and therefore lies closer to them in feature space, while *s*(*x*) is approximately unchanged. As *r*(*x*) is the denominator of the log-ratio in Eq 7, we hypothesised that this would generate positive bias in estimated KL and negative bias in MIND. On the other hand, downsampling is predicted to increase *r*(*x*), because aggregating over multiple source vertices yields fewer target values spanning a comparable range of feature space, while *s*(*x*) again remains comparatively stable. This effect hypothetically produces negative bias in KL and positive bias in MIND.

To investigate these hypotheses in simulated data, we drew samples from Gaussian distributions, independently varying the degree of up- and downsampling applied to each distribution, and computed bias in estimated KL and MIND relative to a ground truth, closed-form solution. As expected, these simulations revealed that upsampling produced positively biased KL divergence values and negatively biased MIND values, reaching a MIND bias of −28% when both distributions were maximally upsampled, while downsampling produced bias of opposite sign, reaching +26% (**Figure 4a-b**).

**Figure 4:**
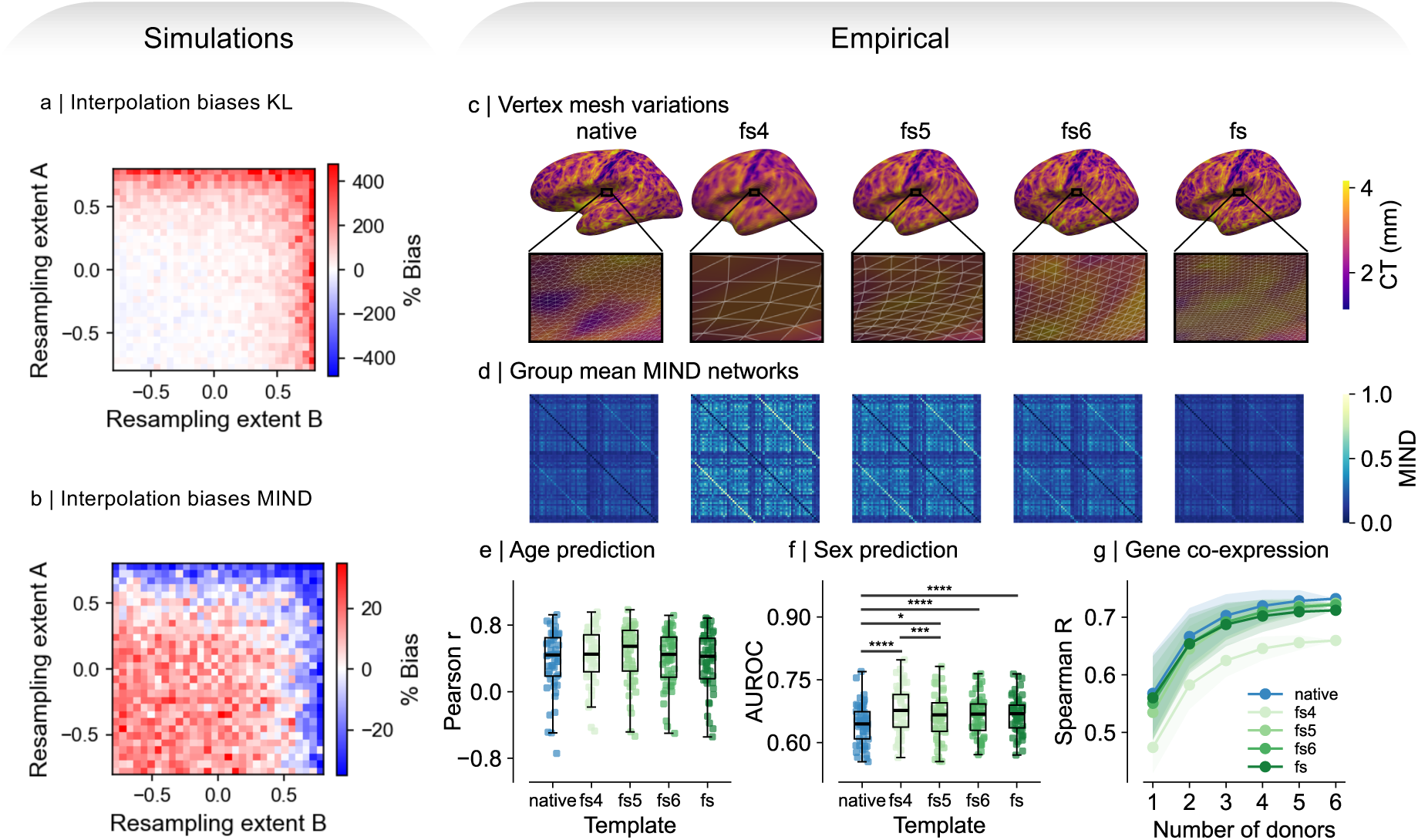
Interpolation biases KL divergence estimation. Percentage bias in **a** symmetric KL divergence (JEF) and **b** MIND as the degree of resampling of each distribution was independently varied. Here, a positive resampling extent indicates upsampling, and negative indicates downsampling. **c** Example cortical thickness maps evaluated in native space (‘native’), fsaverage4 (‘fs4’), fsaverage5 (‘fs5’), fsaverage6 (‘fs6’), and fsaverage (‘fs’) common spaces for ‘bert’, the fully reconstructed adult subject distributed within the FreeSurfer package^53^. **d** Group mean MIND networks estimated using 5 macrostructural features and DK68 parcellation in each analysis space. **e** There were no significant differences in age prediction accuracies between MIND networks generated in the different analysis spaces considered (omnibus Friedman p = 0.10). **f** Sex prediction classification by all common space MIND specifications was more accurate than native space specifications (native vs. fs4: p < 0.001, native vs. fs5: p = 0.0469, native vs. fs6: p < 0.001, native vs. fs: p < 0.001). In **e-f**, bars indicate pairwise significant relationships following the Nemenyi post-hoc test, with **\*** p < 0.05, ** p < 0.01, *** p < 0.001, **** p < 0.0001. **g** Correlations between group mean MIND networks and gene co-expression networks are similar across analysis spaces, except fs4, which was reduced. Lines show the mean and shaded areas show ±1 standard deviation of correlation values across all non-empty combinations of donors used to estimate gene co-expression networks.

#### 3.3.3 Effects of analysis space and vertex mesh resolution on MIND in structural MRI data

Morphological feature maps were computed in native space, containing 97,000-168,000 (mean = 129,000) vertices in the left hemisphere, and in four common spaces: fsaverage, fsaverage6, fsaverage5, and fsaverage4 containing 163,842, 40,962, 10,242, and 2,562 samples per hemisphere, respectively (**Figure 4c**). MIND networks were estimated using these five vertex mesh specifications, in each case using 5 macrostructural features and the DK68 parcellation. In agreement with simulation results, higher resolution templates had overall reduced MIND edge weights (**Figure 4d**).

The biological validity of these networks was compared using age and sex prediction and gene co-expression correlation tasks. We observed no significant differences in age predictions between MIND networks estimated using different analysis spaces (omnibus Friedman p = 0.10; **Figure 4e**), while MIND networks estimated using every common space produced significantly more accurate predictions of sex than native space networks (**Figure 4f**). Notably, we observed that sex classification accuracies were consistently higher when eTIV was not regressed out of edge weights, across all analysis spaces (median AUROC with eTIV regressed = 0.64-0.68, without eTIV regressed = 0.79-0.83), though this was not the case for age predictions (median Pearson r with eTIV regressed = 0.42-0.54, without eTIV regressed = 0.43-0.54, **Supplementary Figure 5**). These results suggest i) that brain volume may mediate some of the effect of biological factors, especially sex, on MIND network edge weights and ii) that this mediation is not solely due to changes in total vertex counts. We did not observe any significant differences in prediction accuracies *between* common space resolutions. MIND networks estimated in all analysis spaces produced similarly strong correlations with gene co-expression, except for fs4, which showed reduced correlations with gene co-expression (**Figure 4g**).

#### 3.3.4 Summary of bias risks and mitigations for analysis space and vertex mesh resolution

In summary, we hypothesised and demonstrated in simulated data that changing vertex mesh resolution using interpolation can bias estimation of KL divergence and MIND. However, we did not observe in empirical data any significant differences in task performance between MIND networks estimated in common space using template meshes of different resolutions, except that the lowest resolution template, fs4, had reduced correspondence with gene co-expression networks. This precludes the recommendation of a single common space template for MIND network analysis.

Furthermore, though native space MIND networks were not outperformed by any common space MIND networks in the age prediction task, they were outperformed by all common space MIND networks in the sex prediction task. We note that UKB adults have low inter-individual variability in total vertex counts, so are subject to relatively uniform interpolation extents when resampled to a common template. For datasets with greater variability in vertex counts, such as the dHCP, subjects would be subject to more variable extents of interpolation when resampled to a common template, theoretically incurring more variable degrees of bias, which could lead to an alternative pattern of results on equivalent biological benchmarks. Finally, regressing eTIV out of MIND edges strongly reduced sex classification accuracies, suggesting that brain size may mediate the effect of sex on MIND similarity.

### 3.4 Parcellation scheme

#### 3.4.1 Sources of variation in parcellation region size

The choice of parcellation scheme used to define brain network nodes is a critical preprocessing decision influencing downstream network properties^75^. Parcellations differ in terms of their granularity, varying from fine- to coarse-grained, and their homogeneity, with homogeneous parcellations containing regions of similar sizes and inhomogeneous parcellations containing regions of different sizes. While the influences of parcellation granularity on downstream neuroimaging analyses have been explored^73,75^, parcellation size homogeneity is not routinely quantified. To address this, we quantified the inhomogeneity in vertex counts of several parcellations using the Gini coefficient. This analysis revealed that the most inhomogeneous atlases were von Economo parcellations, while the most homogeneous was DK318 (**Supplementary Figure 6**).

#### 3.4.2 Effects of parcellation region sizes on KL and MIND estimation in simulated Gaussian distributions

The size of a region influences its morphological feature distributions in two key ways. First, large regions span a greater range of cortical morphology than small regions. For instance, a large region may cover several gyri and sulci, while a small region may cover a single gyrus or sulcus. Larger regions therefore tend to have higher variance in morphological feature values. Second, because larger regions are larger samples of the cortex as a whole, regional means converge to the global cortical mean as regional size increases, reflecting the law of large numbers^76^. We hypothesised that increases in variance and globally convergent mean values of larger regions tend to increase their overlap with other regional distributions, decreasing KL divergence and increasing MIND.

To test this hypothesis, a spatially autocorrelated Gaussian random field was generated across a 2-dimensional grid, non-overlapping regions of independently varied sizes were grown from random seed points, and KL divergence and MIND were estimated between simulated regions. Because points were not drawn from ground truth Gaussian distributions, but were instead defined by region size, absolute KL and MIND were quantified rather than bias relative to an analytic ground truth. Simulations revealed that KL increased and MIND decreased when either region was small, whereas KL decreased and MIND increased when both regions were large (**Figure 5a-b**).

**Figure 5:**
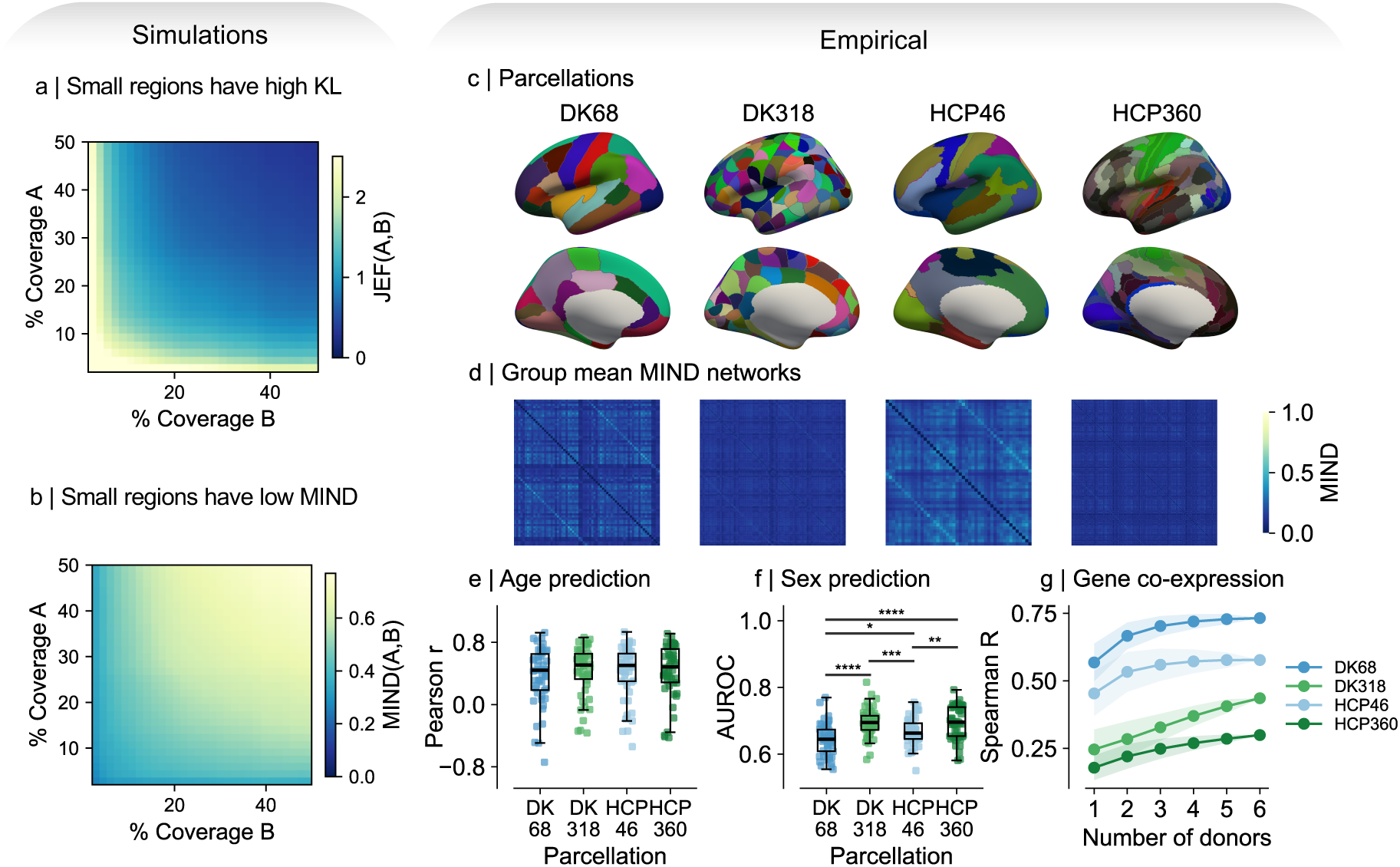
KL divergence and MIND are influenced by parcellation choice. **a** symmetric KL divergence (JEF) and **b** MIND values are influenced by relative region sizes, simulated as the percent coverage of a grid of simulated points. **c** Hierarchically nested parcellations used in empirical data analyses. **d** Group mean MIND networks estimated using parcellations in **c**. **e** Choice of parcellation does not significantly impact the accuracy of age predictions. **f** Finer parcellations produce significantly more accurate predictions of sex (DK68 vs. DK318: p < 0.001, HCP46 vs. HCP360: p = 0.007). In **e-f**, bars indicate pairwise significant relationships following the Nemenyi post-hoc test, with **\*** p < 0.05, ** p < 0.01, *** p < 0.001, **** p < 0.0001. **g** Coarser parcellations had stronger correlations with gene co-expression. Lines show the mean and shaded areas show ±1 standard deviation of correlation values across all non-empty combinations of donors used to estimate gene co-expression networks.

#### 3.4.3 Effects of parcellation region sizes on MIND in structural MRI data

To explore the effects of parcellation granularity on the biological validity of MIND networks, we estimated MIND networks using two pairs of hierarchically nested parcellations, where each pair shared the same gross spatial organisation but differed in granularity (**Methods 2.6.1**, **Figure 5c-d**). We did not directly explore the effects of parcellation homogeneity on MIND network validity in this study as the effects of homogeneity are difficult to isolate from genuine differences in biology between parcellations.

Fine-grained parcellations had higher edge-level variability, reflecting estimation of similarities between smaller and noisier samples of cortex (**Supplementary Figure 7**). In line with simulations, we observed that regional MIND weighted degree was correlated with region size, except for the DK318 parcellation (**Supplementary Figure 8a**). We hypothesised that the reduced association between region size and weighted degree in DK318 could reflect the reduced variability in region sizes of this parcellation (**Supplementary Figure 6**). To test this, we assessed the correlation between the region size-weighted degree correlation value and size homogeneity, indexed using the Gini coefficient, of 17 parcellations, and observed a strongly significant correlation (Pearson r = 0.70, p = 0.002, **Supplementary Figure 8b**).

We observed no significant differences in the accuracies of age predictions between parcellation schemes (omnibus Friedman p = 0.10, **Figure 5e**), though finer parcellations produced significantly more accurate predictions of sex (**Figure 5f**). Finer parcellations were less strongly correlated with matched gene co-expression networks (**Figure 5g**), potentially reflecting the noisier estimation of edges between smaller parcels, though correlations could also be influenced by differences in the number of edges used. Additional analyses revealed that coarse graining before MIND estimation, that is, computing similarities between coarse-grained parcels, rather than averaging across edges of a MIND network estimated using a fine-grained parcellation, were associated with improved predictions of age and sex, though non-significantly, and improved correspondence with gene co-expression networks (**Supplementary Figure 9**).

#### 3.4.4 Summary of effects of parcellation choice on KL divergence and MIND

In summary, theoretical, simulated, and empirical analyses revealed that KL and MIND values are influenced by the size of parcellated regions. Larger regions, by virtue of sampling more of the cortex, are more similar to the rest of the brain on average, though we observed that this coupling was weaker for more homogeneous parcellations. This suggests that conducting sensitivity analyses using a parcellation of differing homogeneity could be a useful approach to assess whether MIND network associations are driven by regional volume.

Finally, we did not find evidence that fine-grained or coarse-grained parcellations produced MIND networks of systematically enhanced biological validity: the appropriateness of the parcellation depends on the task at hand and so remains at the discretion of the investigator. However, we observed that coarse graining *prior* to network estimation, i.e. computing pairwise similarities between coarse parcels, produced MIND networks with favourable performance to networks that were coarse grained *after* MIND estimation by averaging across fine grained parcels.

### 3.5 Feature covariance

#### 3.5.1 Sources of feature covariance

Selection of optimal features for estimation of MIND networks should primarily be performed based on relevance to the scientific question and can additionally be informed by vertex-level interpretability of features, maximisation of heritability, and reducing redundancy^18^. An additional property that influences the technical validity of multivariate MIND networks is covariance between features, which may be genetically driven. For example, genetic analyses of cortical structural phenotypes in UKB data identified four latent structural factors relating to cortical expansion, cortical curvature, neurite density and orientation, and water diffusion, with phenotypes loading onto the same factor tending to be highly correlated at the genetic and phenotypic levels^54^.

#### 3.5.2 Effects of feature covariance on KL and MIND estimation in simulated Gaussians

The inclusion of several highly covarying features indexing the same underlying construct weights the estimated KL divergence towards similarities in the over-represented construct. Furthermore, covariance can directly bias the k-NN estimator. As noted, the k-NN estimator computes nearest-neighbour distances within a *d*-dimensional ball assumed to enclose a locally uniform probability density (**Methods 2.3.1**). However, covariance elongates the probability density along the covarying dimensions, and differences in covariance between distributions lead to an area of probability density being densely sampled by one distribution and sparsely by the other. This can generate local non-uniformity within the *d*-dimensional ball, biasing the k-NN estimator (**Methods 2.3.1**). We therefore hypothesised that differences in covariance between distributions would bias KL divergence and MIND.

To assess the impact of feature covariance on KL divergence and MIND, we drew samples from multivariate Gaussian distributions, independently varying the degree of covariance between dimensions, and computed bias in estimated KL and MIND relative to a ground truth, closed-form solution. These simulations demonstrated that higher covariance between features resulted in inflated k-NN estimated KL divergence values and suppressed MIND values, with a maximum absolute MIND bias of −62% observed when there were large differences in covariance between the simulated regions, in line with our prior hypothesis (**Figure 6a-b**).

**Figure 6:**
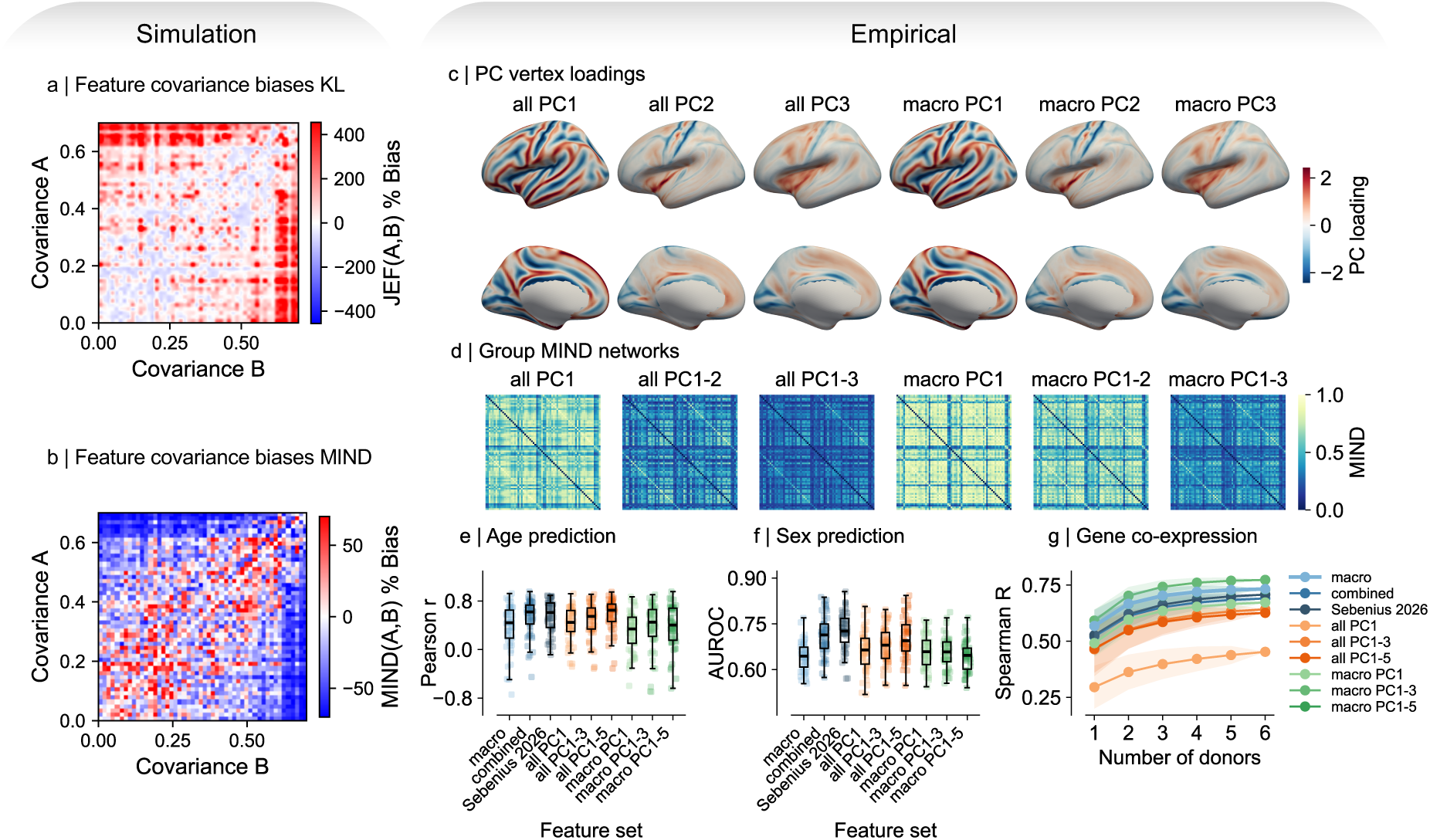
Covariance biases KL divergence estimation and principal components are suitable input features to MIND. Percentage bias in **a** symmetric KL divergence (JEF) and **b** MIND as the covariance between features within each distribution was independently varied. **c** Vertex loadings on PC1, PC2, and PC3, computed from all 10 structural features (‘all’) or from 5 macrostructural features only (‘macro’), for group mean feature maps. **d** Group mean MIND networks estimated using combinations of PC1-3 computed from all structural features or 5 macrostructural features. In the following plots, blue colours signify MIND networks estimated using combinations of raw features, orange colours signify MIND networks estimated using combinations of principal components computed on all 10 macro- and microstructural features (‘all’), and green colours signify MIND networks estimated using combinations of principal components computed on 5 macrostructural features only (‘macro’). Pairwise significance bars are not shown due to space constraints. **e** Most accurate age predictions were obtained when using all_PC1-5 (median Pearson r = 0.65), which reached significance when compared to the default feature set (p = 0.004), but not when compared to the ‘combined’ feature set (p = 1.00). **f** Most accurate predictions of sex were obtained when using the ‘Sebenius 2026’ feature combination (median AUROC = 0.73), which reached significance when compared to networks generated using all combinations of PCs, except all_PC1-4 (p = 0.09) and all_PC1-5 (p = 0.76). **g** Highest correlations with gene co-expression were obtained when using PC1-3 computed on macrostructural features (Pearson r = 0.77 when N = 6 donors used). Lines show the mean and shaded areas show ±1 standard deviation of correlation values across all non-empty combinations of donors used to estimate gene co-expression networks.

#### 3.5.3 Effects of feature covariance on MIND in structural MRI data

Based on theoretical understanding and simulation results that KL divergence is biased by feature covariance, we orthogonalised input MRI features using principal component analysis of vertex maps (**Methods 2.7.4**), and estimated MIND networks from combinations of the resulting PCs. Feature sets used in computation of principal components (PCs) were either all 10 macrostructural and microstructural features (‘all’) or 5 macrostructural features only (‘macro’). MIND networks were generated using all combinations of PC1-3, as well as PC1-4, and PC1-5 using both the ‘all’ and ‘macro’ PC sets. We additionally generated MIND networks from four combinations of raw features: 5 macrostructural features (‘macro’); 5 microstructural features (‘micro’); all 10 macro- and microstructural features (‘combined’); and a set of four features selected on the basis of interpretability, heritability, and non-redundancy: cortical thickness, mean curvature, mean diffusivity, and intracellular volume fraction (‘Sebenius 2026’)^18^. All networks were computed in native space using the DK68 parcellation. In total, we estimated MIND networks using 22 feature combinations and compared the validity of these permutations on age and sex prediction and gene co-expression correlation tasks. Performance for a subset of 9 configurations is visualised in **Figure 6**, and the full set is visualised in **Supplementary Figure 13**.

For PCs computed on all features, the first four components each explained ≥10% of the total variance, collectively explaining 76.6% of variance in cortical morphology (**Supplementary Figure 10a**); whereas for PCs computed on macrostructural features only, the first three components explained ≥10% of the total variance, collectively explaining 92.7% of variance in cortical morphology (**Supplementary Figure 11a**). Vertex and MRI feature loadings on all components are shown in **Supplementary Figures 10-11**. Component loadings were highly stable to subject-level subsampling within the UKB dataset (median congruence across 100 subsamples ≥0.91 for all components; **Supplementary Figure 10c**, **11c**) and were congruent with loadings derived externally from the dHCP dataset (**Supplementary Figure 12**).

Vertex loadings on PC1-3 for both input feature sets are shown in **Figure 6c**, and MIND networks computed from various combinations of PCs are shown in **Figure 6d**. MIND networks estimated using ‘all PC1-5’ produced the most accurate predictions of age, and this reached significance when compared with the default macrostructural feature set, but not when compared to networks estimated from combinations of macro- and microstructural features (**Figure 6e**). MIND networks estimated using ‘Sebenius 2026’ produced the most accurate predictions of sex, and this reached significance when compared to networks estimated using every combination of PCs, except for PC1-4 and PC1-5 (**Figure 6f**). Macrostructural-only PC1-3 had the highest correlations with gene co-expression (**Figure 6g**).

#### 3.5.4 Summary of bias risks and mitigations for feature covariance

In summary, theoretical and simulated data analyses support the conclusion that feature covariance can introduce bias into the k-NN estimator when distributions display differences in covariance. However, in empirical data, we did not observe systematically improved task performance for MIND networks computed using principal components as input features. Therefore, while PCA remains an option for obtaining a set of non-redundant features with reduced regional covariance, feature selection ultimately depends on the biological question at hand.

## 4 Discussion

Using simulations and empirical data, we investigated the effects of five properties of input feature maps on MIND network estimation: smoothness, presence of identical values, analysis in native or common space and vertex mesh resolution, parcellation choice, and feature covariance. We investigated the biological validity of MIND networks estimated from healthy UK Biobank adults using a range of preprocessing configurations via age and sex prediction tasks and correlations with gene co-expression and largely confirmed our primary hypothesis that feature distribution properties introducing bias in simulations, when evident in empirical feature maps, led to sub-optimal MIND networks with reduced prediction accuracies and correlations with gene co-expression. We discuss each of these properties in turn, present a set of recommendations for researchers wishing to use MIND in their own work, and conclude with limitations and outstanding questions.

### 4.1 Discussion of bias sources

#### 4.1.1 Smoothing

Smoothing is applied in typical neuroimaging preprocessing pipelines for several reasons, though primarily to enhance signal to noise ratio^72,73^. We observe, however, that increases in vertex map smoothing may *decrease* the signal to noise ratio in MIND networks, evidenced by increases in the variability of edges across subjects, decreased accuracy of sex predictions, and decreased associations with gene co-expression. Previous work has also cautioned against excessive smoothing: smoothing reduces morphometricity, the proportion of variance in diverse traits explained by vertex-wise morphometrics^77^; can reduce age prediction accuracy by flattening meaningful between-subject variability^73^; and blurs areal boundaries, leading to reduced spatial concordance of parcellated label assignments across subjects^78^.

#### 4.1.2 Identical values

Identical values can be associated with normal biology, as we show is the case in the smooth cortices of early neonatal brains, or can be inadvertently introduced during routine preprocessing pipelines, for example due to quantisation following storage of data in efficient but imprecise data formats, or due to upsampling using nearest neighbours interpolation. We observed that a high proportion of identical values, introduced due to quantisation, generated bias in KL divergence and MIND estimated in simulated data, and reduced the biological validity of MIND networks.

#### 4.1.3 Analysis space and vertex mesh resolution

Morphometric feature maps can be generated in native or common space, with native space typically preferred to avoid geometric distortion introduced by warping to a template^79^. However, we demonstrated that native space MIND networks generally had reduced prediction accuracies relative to common space MIND networks. Native space analysis permits inter-individual variability in total and regional vertex counts, thus permitting inter-individual variability in the degree to which regional morphometrics are appropriately sampled, and generating individual-specific patterns of bias, or noise, that may reduce the accuracies of edge-based prediction models.

We observed that interpolation-based up- or downsampling generated bias in KL and MIND estimated in simulated data. When there is low inter-individual variation in total vertex counts, as in the adult UKB cohort, overall bias due to interpolation incurred by warping to a common template may be acceptably small. However, in datasets with high inter-individual variation in total vertex counts, as in the neonatal dHCP cohort, subjects require vastly differing degrees of up- or downsampling, therefore introducing differing degrees of interpolation-induced bias, which may make common space analysis less suitable. This highlights a potential trade-off between bias brought about by: i) inter-individual variability in vertex counts and thus in appropriateness of sampling of regional morphometrics when using native space analysis; and ii) interpolation-based resampling when using common space analysis.

If analysing data in common space, our results suggest that performance may be improved by selecting lower-resolution common space templates, though this trend did not reach significance. We therefore suggest fsaverage6 as a reasonable default, on the basis that it retains sufficient regional sample counts to support fine-grained parcellations without incurring the additional interpolation required to reach the highest-resolution template.

Furthermore, we observed that regressing edge weights on eTIV reduced sex prediction accuracies, and that this effect was present in native *and* common space analyses, suggesting that brain volume mediates some of the effect of sex on MIND network edge weights *independently* of differences in total sample counts. In line with previous neuroimaging recommendations^48^, brain volume may therefore be a useful covariate to consider in statistical models when the goal is to identify biological associations with MIND networks above and beyond those mediated by brain volume.

#### 4.1.4 Parcellation scheme

Parcellation choice is a long-standing consideration in network neuroscience, with node size influencing graph-theoretic metrics derived from diffusion MRI-based networks for example^75^. We observed in simulated and empirical data that larger parcels tended to have lower KL and higher MIND values as: i) they cover a larger range of morphologies, increasing their variance and thus their overlap with other regions and ii) their means converge to the mean over the whole cortex by the law of large numbers, bringing their means closer, on average, to other distributions. Though we indexed region size using vertex counts, surface area and volume are also measures of region size, so are expected to be correlated with weighted degree under some conditions.

Importantly, the strength of this association was modulated by the size homogeneity of the parcellation, being attenuated in homogeneous parcellations. Investigators using MIND in their own research may therefore benefit from performing sensitivity analyses using parcellations of differing homogeneities to assess whether associations with MIND are driven by regional volumes. This approach may be preferable to controlling for regional volume in analyses of MIND phenotypes when MIND networks use volume as an input feature, as this would be directly controlling for a mediator of interest^80^.

Finally, though previous studies using mean regional morphometrics have found that finer parcellations have improved age prediction accuracies^73^ and carry higher morphometricity^81^, we did not find any evidence that performance of MIND networks was systematically improved by parcellations of a particular grain. Parcellation choice should ultimately follow from the biological question at hand.

#### 4.1.5 Feature covariance

MIND was initially validated using the default five macrostructural features generated by FreeSurfer, minimising parameter selection, maximising reproducibility, and lowering the barrier to adoption for researchers without extensive computational experience. However, we observed that alternative feature sets may produce favourable performance in different biological tasks, highlighting that feature selection should primarily be performed in relation to the scientific question. An example of question-oriented feature selection can be found in Sebenius et al.^18^, who selected features with high heritability to estimate MIND networks for use in downstream genetic analyses. These analyses identified gradients of genetic patterning of morphology that recapitulated long-standing hypotheses as to the phylogenetic origins of cytoarchitectonic variation in the cortex^17^.

We observed that feature covariance generated bias in KL divergence and MIND in simulated data, motivating the use of principal component analysis to generate statistically orthogonal feature sets for use in MIND network estimation. The first principal component largely reflected sulco-gyral differences, in line with this being a major axis of cortical morphological variation^82–84^, and was externally replicable between the UKB and dHCP datasets. Beyond removing redundancy, these components capture latent modes of cortical organisation rather than measurements that happen to be available, and so may align more closely with the biology underlying morphometric similarity^85^. Orthogonalising the feature space therefore shifts MIND away from being constrained by what is conveniently measured and toward a more data-driven representation of cortical structure^86^. However, MIND networks estimated from combinations of principal components did not provide systematically favourable task performance in empirical data, indicating that while PCA can be used to obtain a non-redundant feature set with attenuated covariance, the ‘optimal’ combination of input features depends on the biological task or question at hand.

### 4.2 Recommendations to researchers

Based on our findings, we make several recommendations to researchers wishing to use MIND in their own work (**Table 3**).

**Table 3:** Recommendations to researchers. The resample=True flag is supported in the current MIND repository^87^. An updated version of the repository will support diagnostic outputs, including global and regional proportions of identical values, and computation of principal components for use as input features.

| Pipeline step or bias source | Recommendations |
| --- | --- |
| Smoothing | Avoid smoothing feature maps prior to MIND network estimation where possible. |
| Identical values | Avoid storing input data in imprecise formats, or upsampling using nearest neighbours interpolation, as these procedures can introduce identical values.<br>If computing MIND from a single feature, check that the global percentage of identical values is below 90%. If it is >90%, run univariate MIND using the `resample=True` flag. |
| Analysis space | If the inter-individual variability in brain size is low, consider analysing data in common space. If inter-individual variability in brain size is high, native space analysis may be preferable.<br>If using a common space template, we recommend `fsaverage6` as a default, which balances biological validity with sufficient regional sample counts to support fine-grained parcellations.<br>Consider including total brain volume as a covariate in statistical models. |
| Parcellation selection | Parcellation selection should be performed primarily based on the scientific question.<br>Consider conducting sensitivity analyses using a parcellation of a different grain.<br>If using a coarse-grain parcellation, coarse-graining should be performed prior to network estimation where possible. |
| Feature selection | Feature selection should be performed primarily based on the scientific question, or based on maximising interpretability, non-redundancy, or heritability<sup>18</sup>.<br>Vertex-wise principal component loadings can be used to ensure that features capture non-redundant information. |

### 4.3 Limitations and future work

Our study has limitations. Our simulations used Gaussian distributions so that the closed form solution for KL divergence could serve as a ground truth for estimation of bias in KL divergence and MIND. However, regional vertex-wise morphological feature distributions often diverge from normality, demonstrating high skew^88,89^ or bimodality, given that sulco-gyral patterning is a strong organiser of cortical organisation^82–84^ and regions can span multiple gyri and sulci. Future work may therefore benefit from simulating regional feature distributions as non-Gaussian, for example using more skewed distribution shapes, such as the beta or gamma distributions, for which closed-form solutions exist^46^.

We assumed that MIND networks with higher overall bias, predicted theoretically and via simulations, would be less valid measures of coordinated patterns of brain morphology, which we proxied using predictions of age and sex and correlations with gene co-expression networks. Cross-validated predictions of age and sex are widely-used benchmarks of MIND networks^4,11–13^ and neuroimaging phenotypes more broadly^73,90,91^ given the sensitivity of brain structure to these overt biological characteristics. Furthermore, we used correlations with gene co-expression as a biological validation given that previous work has observed strong correlations between structural similarity and gene co-expression^4,7,13^, and it is thought that similarity at the macroscale reflects underlying genetic or transcriptional similarity^2^, reflecting Cheverud’s conjecture^50^. However, gene co-expression does not constitute a definitive ‘gold standard’ against which to validate structural similarity networks due to the influence of non-transcriptomic factors on brain structure, such as experience-dependent plasticity^92^. Future work could extend our biological benchmarking framework by: i) predicting additional phenotypes, such as cognition, given strong evidence for the brain’s network architecture being a substrate for underlying cognition^93–95^; ii) using statistical approaches beyond multivariate prediction, such as univariate associations^73^ or morphometricity^77,96^; iii) validating networks using genetic approaches, such as twin-based heritability^4^; and iv) benchmarking a greater number of alternative combinations of preprocessing configurations.

Finally, the core aim of this study was to develop and share an optimal pipeline for MIND estimation without changing the MIND estimation algorithm itself. Future work could benchmark alternative estimators of KL divergence, for example with in-built bias mitigations^97^, alternative information theoretic measures of divergence, such as the Jensen-Shannon divergence^98–100^, or alternative normalisations of these divergence measures to generate similarity networks.

## 5 Conclusion

We investigated sources of bias in estimation of MIND networks using simulated and real-world structural MRI data and largely found that sample properties shown to bias KL divergence and MIND in simulated data, when introduced to input morphological feature maps, diminished the biological validity of MIND networks. We presented a set of recommendations for researchers wishing to use MIND in their own research and facilitate the adoption of these recommendations via changes to the original MIND GitHub repository^87^, which will be implemented upon full publication of this work.

## 6 Declarations

### 6.1 Data and code availability

All data analysed in this study are publicly available. Code required to reproduce all analyses and results reported in this paper will be made available upon publication at https://github.com/edhutch1/MIND_methods.

### 6.2 Author contributions

A.D., E.D.H., B.C., M.H.G., E.T.B., and S.E.M. conceived of the primary methodology. A.D. performed the simulation analyses and analyses of dHCP data. E.D.H. performed simulation analyses investigating the effects of smoothing and resampling to remove identical values, and analyses of UKB data. B.C., M.H.G., S.K.C., I.S., R.A.I.B., E.T.B., and S.E.M. advised on primary analyses. I.S. developed the methodology for resampling to remove identical values. S.K.C. developed the parallel optimization for the MIND computation. A.D. and E.D.H. drafted the primary manuscript. All authors reviewed and edited the manuscript.

### 6.4 Ethics oversight

All data analysed in this study were previously published and collected in accordance with relevant ethical standards.

### 6.5 Funding acknowledgements

A.D was supported by the EPSRC Research Council, part of the EPSRC DTP (EP/W524475/1). E.D.H was supported by a Rosetrees Trust grant MB2023\100002 and funding from the University of Cambridge School of Clinical Medicine. BC was supported by the Cambridge School of Clinical Medicine Doctoral Training Programme in Medical Research and Pinsent-Darwin Studentship in Mental Pathology. M.H.G. was supported by the National Institutes of Health Intramural Research Program, the NIH Oxford-Cambridge Scholars program, and the Cambridge Trust. I.S. was supported by Samvid Philanthropies and the Gates-Cambridge Trust. E.T.B. was supported by ImmunoMIND, funded by the Medical Research Council (MR/Z50354X/1) as part of the UKRI Mental Health Platform. S.E.M. was supported by a UKRI Medical Research Council Future Leaders Fellowship [UKRI3090]. MRI data were curated and analysed using a computational facility funded by an MRC research infrastructure award (MR/M009041/1) to the School of Clinical Medicine, University of Cambridge and supported by the mental health theme of the NIHR Cambridge Biomedical Research Centre (BRC).

This work was additionally supported by the NIHR Cambridge Biomedical Research Centre (NIHR203312) and the NIHR Applied Research Collaboration East of England. The views expressed are those of the authors and not necessarily those of the NIHR or the Department of Health and Social Care.

