## Supplementary Materials for "MIND the gap: methodological considerations and guidance for structural MRI similarity network analysis with MIND"

### Table of contents

**Supplementary Figures** **1**

1. Impacts of smoothing on feature maps and MIND networks 2
2. Resampling removes bias due to identical values 3
3. Impacts of identical values on MIND networks 4
4. Age-related trends in vertex counts 5
5. Regressing eTIV from MIND edges reduces predictive performance 6
6. Region size inhomogeneity in commonly used parcellations 7
7. Impact of parcellation choice on MIND edge variability 8
8. Parcellation homogeneity modulates the association between region size and degree 9
9. Coarse graining before MIND estimation is optimal 10
10. Principal component analysis of 10 macro- and microstructural features 11
11. Principal component analysis of 5 macrostructural features 12
12. Feature loadings on principal components are externally replicable 13
13. Biological validation of all feature sets 14

**References** **15**

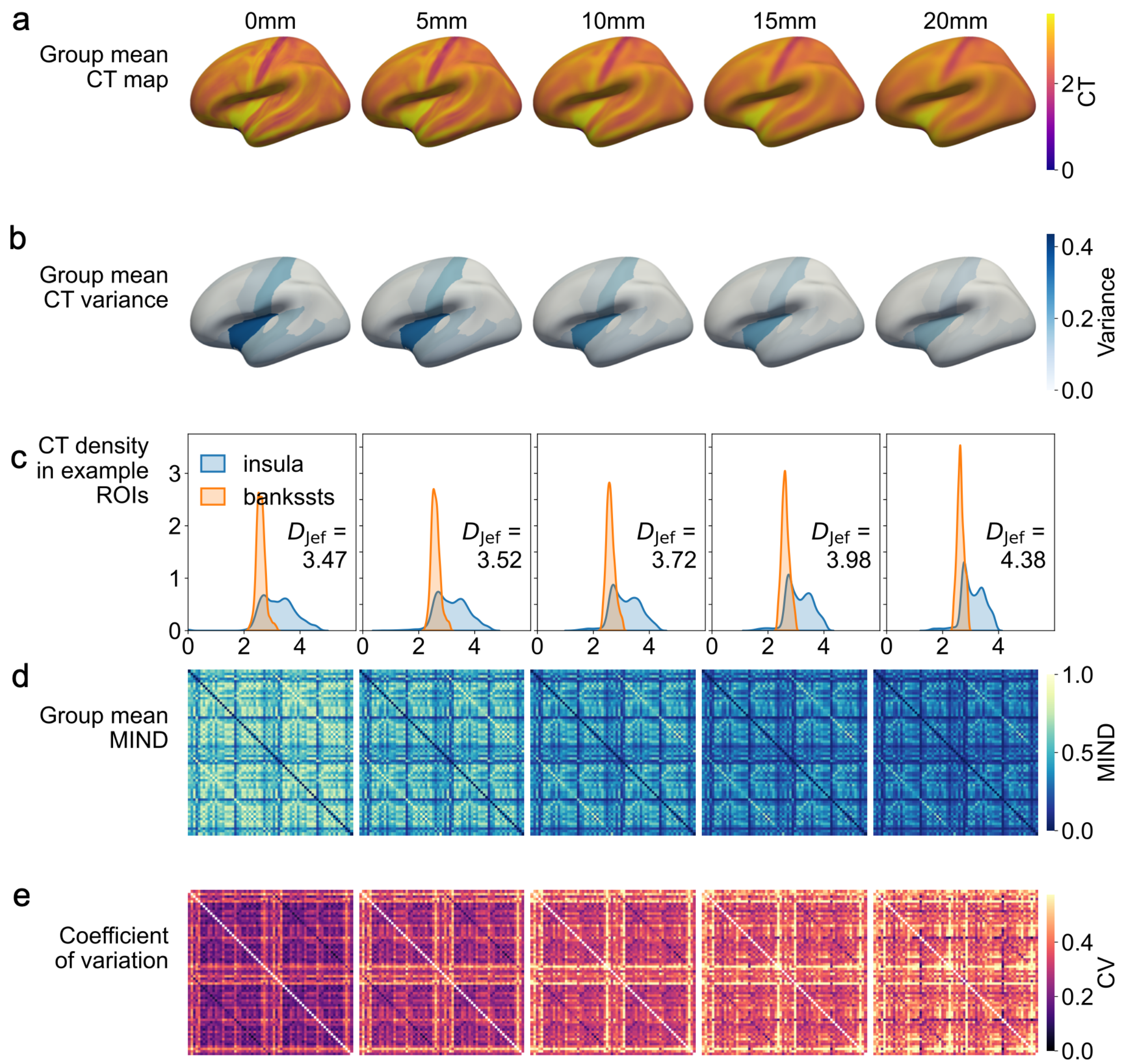


***Supplementary Figure 1: Impacts of smoothing on feature maps and MIND networks.*** *Results in a column correspond to the smoothing level indicated at the top of the column.* ***a*** *Group mean cortical thickness (CT) map smoothed using a Gaussian kernel with FWHM = [0, 5, 10, 15, 20], with 0 mm indicating no smoothing.* ***b*** *Regional CT variances decrease as smoothing increases, shown for the CT maps in* ***a****.* ***c*** *Regional CT distributions show decreases in overlap and increases in KL as smoothing increases, shown for an example pair of cortical regions in the maps in* ***a****.* ***d*** *Group mean MIND networks show decreases in edge weights as smoothing is increased.* ***e*** *Edge-wise coefficients of variation, calculated as the standard deviation/mean of an edge across subjects, increase as smoothing increases.*

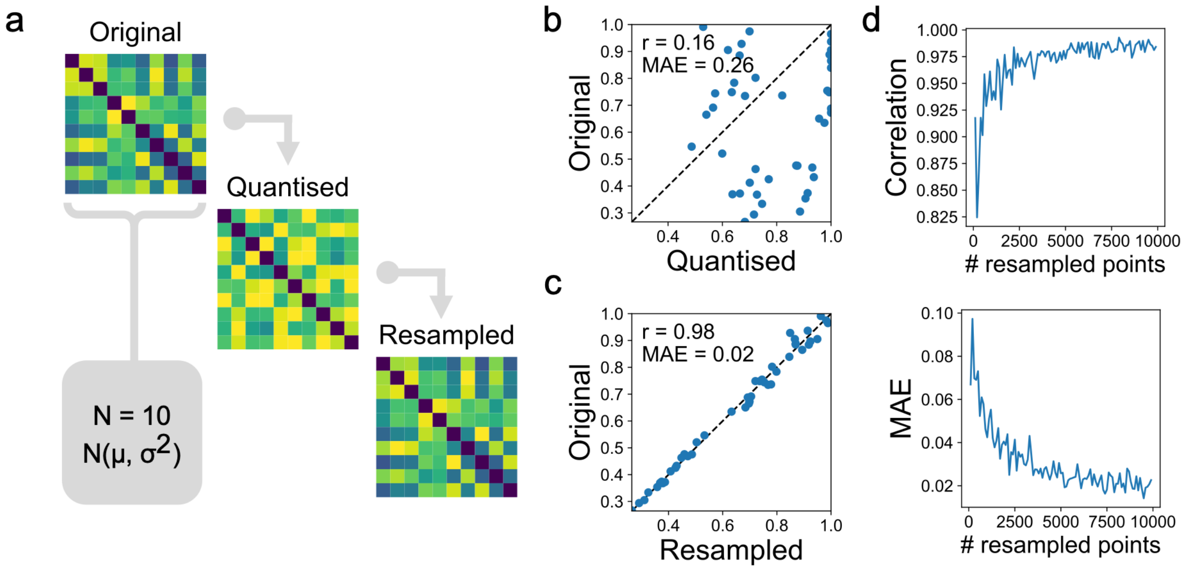


***Supplementary Figure 2: Resampling removes bias due to identical values***. ***a*** *10 gaussian distributions were randomly initialised with mean and standard deviations chosen from* $\mu \in [1.7, 2.9], \sigma \in [0.4, 1.4]$*, reflecting empirical ranges of regional means and variances from the cortical thickness map of a representative adult brain. 10,000 samples were drawn from each distribution, and a MIND network was estimated ('Original'). A random degree of quantisation selected from* $Q \in [10^{-4}, 10^{-2}]$ *was applied to each distribution to introduce identical values, and MIND was recomputed without ('Quantised') and with ('Resampled') resampling.* ***b*** *MIND networks computed from original and quantised data were poorly correlated at the edge level (Spearman r = 0.16, mean edge-wise MAE = 0.26).* ***c*** *Original and Resampled MIND networks were highly correlated at the edge level (Pearson r = 0.98, mean edge-wise MAE = 0.02).* ***d*** *The analysis in* ***c*** *was repeated while drawing a variable number of samples from each region. Agreement between Original and Resampled networks improved with increasing N, plateauing at approximately 4000 samples.*

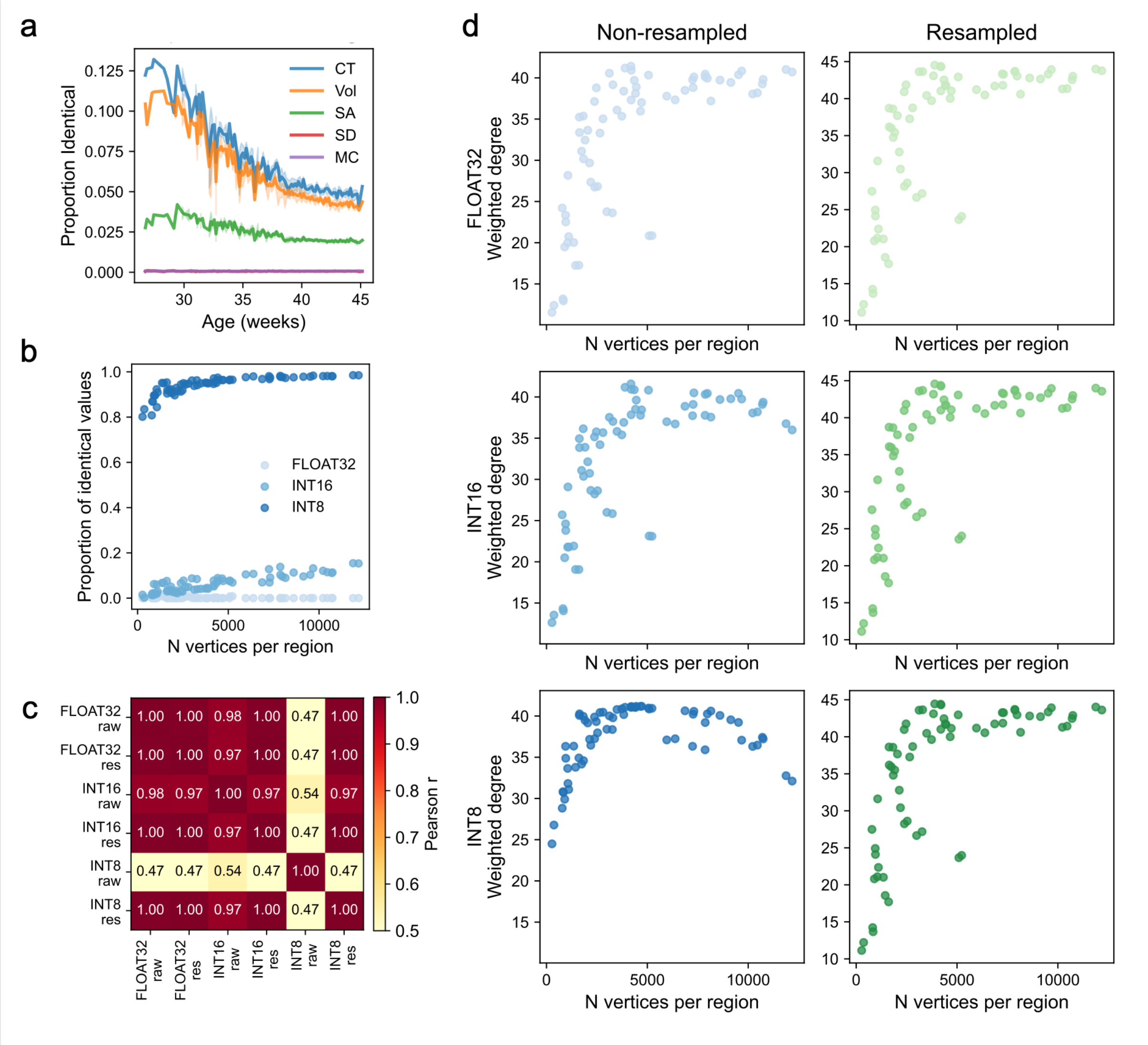


***Supplementary Figure 3: Impacts of identical values on MIND networks**. **a** Mean cortical proportions of identical values over postmenstrual age in the dHCP cohort for five macrostructural features. Panels **b-d** show results from a variably quantised group mean sulcal depth (SD) map, obtained by averaging across individual SD maps in fsaverage space. In each case, maps are parcellated using DK68. **b** Larger regions contain a higher proportion of identical vertex values. **c** Pearson correlations between edges of MIND networks computed from the group mean sulcal depth map quantised to varying extents (FLOAT32, INT16, INT8), and either non-resampled ('raw') or resampled with N = 4000 samples ('res'). **d** Scatterplots showing the relationship between the number of vertices in a region and the MIND weighted degree of that region, for each quantisation condition. Non-resampled INT8 MIND networks show a tight non-linear relationship between number of samples and weighted degree.*

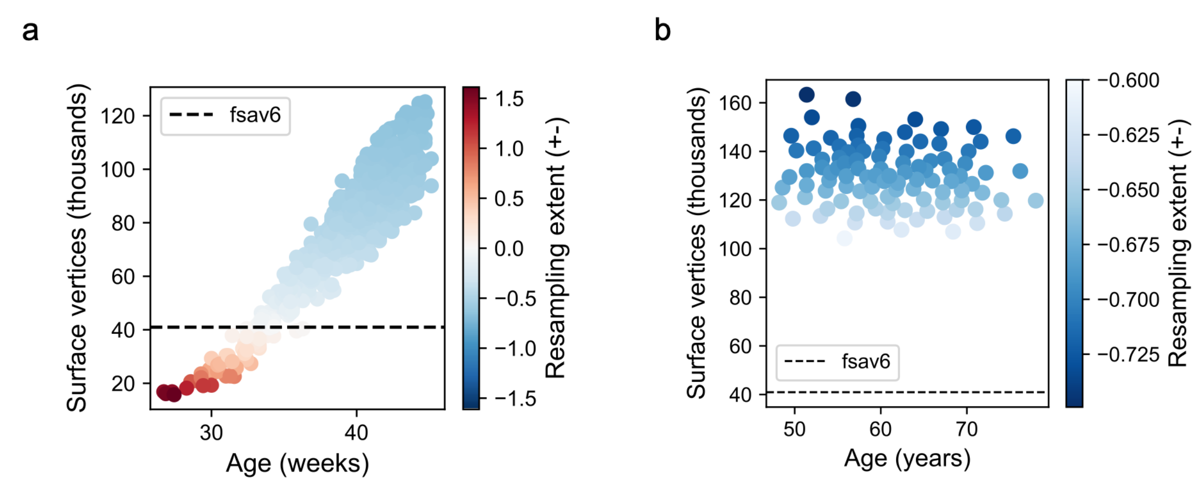


***Supplementary Figure 4: Age-related trends in vertex counts. a*** *Total sample count increases over age in the dHCP cohort. Resampling to fsaverage6 space variably up- and downsamples individual vertex meshes.* ***b*** *Total sample count remains relatively stable in the UKB cohort and resampling to fsaverage6 space more uniformly downsamples individual vertex meshes. For the purposes of visualising UKB data, individual data points were micro-aggregated into groups of ≥5 using the maximum distance to average vector algorithm[101] and the colour bar range was compressed. In both panels, resampling extent was computed as (template - native)/native sample counts.*

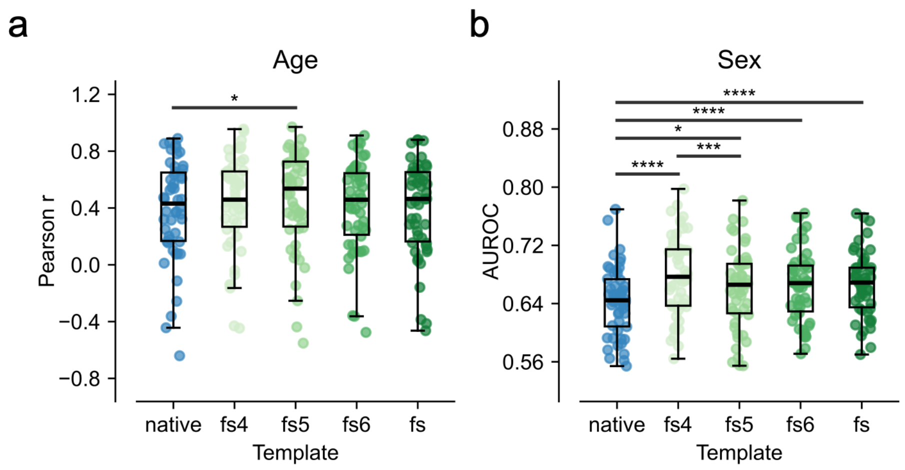


| | native | fs4 | fs5 | fs6 | fs |
|---|---|---|---|---|---|
| eTIV | **0.441** | 0.451 | **0.544** | 0.450 | 0.425 |
| -eTIV | 0.431 | **0.459** | 0.537 | **0.458** | **0.464** |

| | native | fs4 | fs5 | fs6 | fs |
|---|---|---|---|---|---|
| eTIV | 0.644 | 0.677 | 0.666 | 0.668 | 0.669 |
| -eTIV | **0.803** | **0.785** | **0.803** | **0.821** | **0.825** |

***Supplementary Figure 5: Regressing eTIV from MIND edges reduces predictive performance. a** Age and **b** sex prediction results for MIND networks estimated in each analysis space, shown here without regression of eTIV from MIND edge weights prior to model training. Beneath each plot, median performance across folds (Pearson r for age, AUROC for sex) is given both with ('eTIV') and without ('-eTIV') regression of eTIV, with the higher of the two in bold. When eTIV is regressed, age prediction is largely unchanged, though sex classification accuracies show systematic and large decreases.*

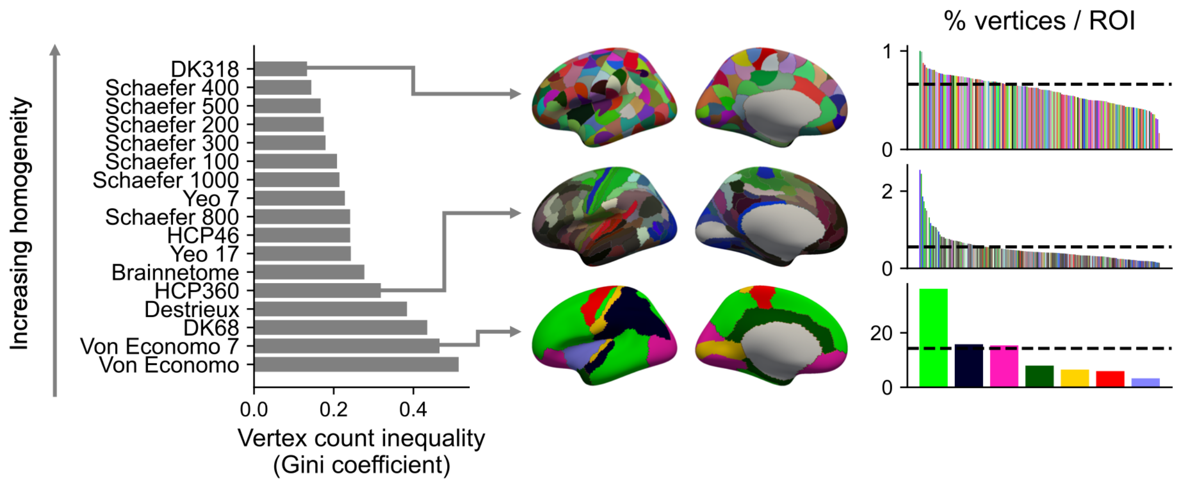


***Supplementary Figure 6: Region size inhomogeneity in commonly used parcellations.*** *Homogeneity in regional vertex counts is quantified using the Gini coefficient for 17 commonly-used parcellations[58–60,102–106] in fsaverage space. For three parcellations of differing homogeneities, ranked left hemisphere regional vertex counts are displayed, with the horizontal dashed line indicating the mean number of vertices.*

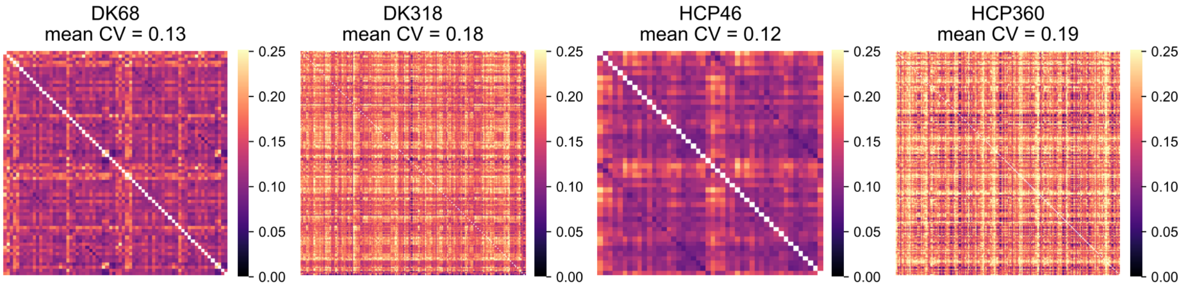


***Supplementary Figure 7: Impact of parcellation choice on MIND edge variability.*** *Edge-wise coefficient of variation, computed as standard deviation/mean across subjects, is globally reduced in coarser parcellations.*

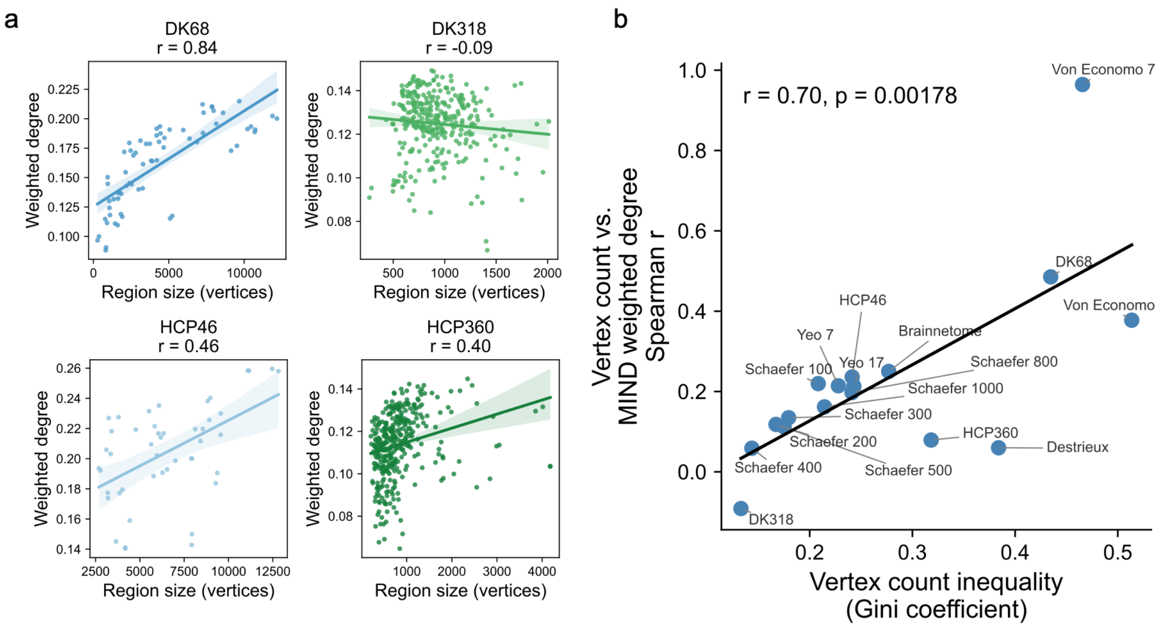


***Supplementary Figure 8: Parcellation homogeneity modulates the association between region size and degree. a** Scatterplots show the relationships between regional MIND weighted degree, computed from group mean MIND networks, and vertex counts, from the fsaverage template, for the four main parcellations considered in this study. Vertex count is moderately or strongly correlated (Spearman r) with weighted degree in all cases except DK318. **b** Across 17 parcellations, the Spearman r value between regional vertex count and MIND weighted degree was significantly correlated with vertex count inhomogeneity (Pearson r = 0.70, p = 0.002). In each case, a single MIND network was computed using 5 macrostructural features, where each feature was the average left hemisphere map computed across all subjects.*

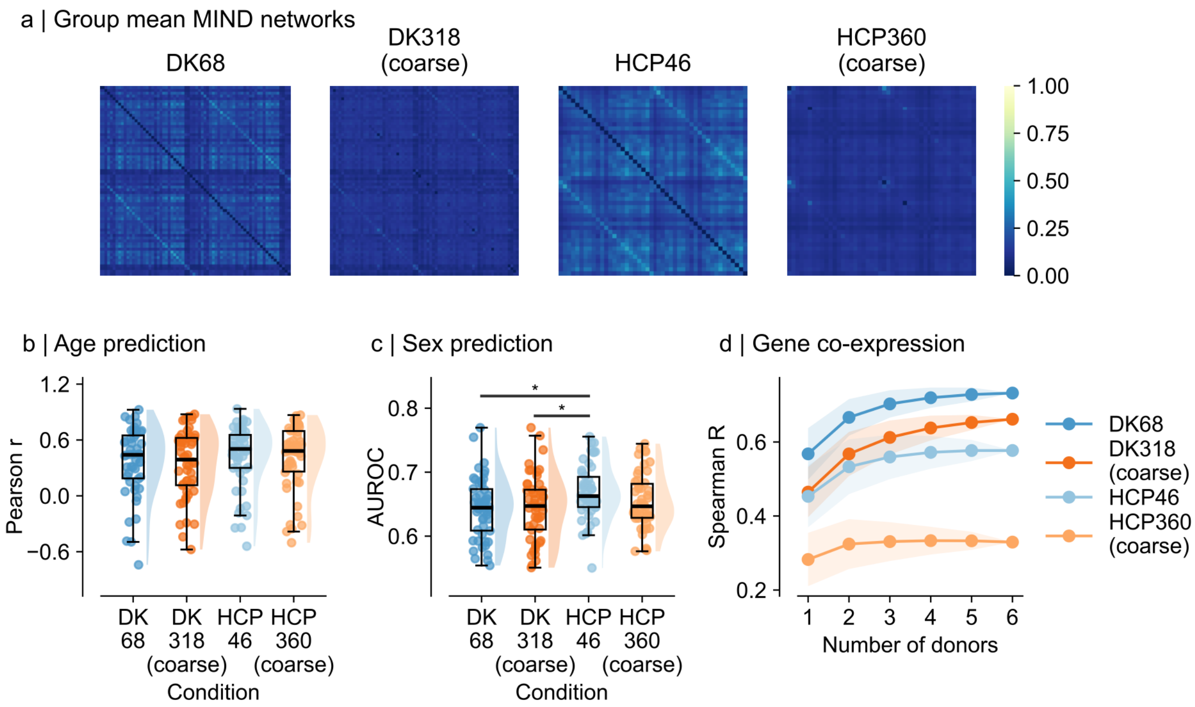


***Supplementary Figure 9: Coarse graining before MIND estimation is optimal. a*** *Coarse-grained group mean MIND networks. For DK68 and HCP46, coarse-graining was performed before MIND estimation, i.e. pairwise similarities were estimated between large, coarse-grained regions, and for DK318 (coarse) and HCP360 (coarse), coarse-graining was performed after MIND estimation, i.e. pairwise similarities were estimated between fine-grained regions, after which edge weights were averaged to obtain the similarities between large, coarse-grained regions.* ***b-c*** *Coarse-graining after network estimation, relative to before, generally led to poorer predictions of age and sex, though these trends did not reach significance. In* ***b-c****, bars indicate pairwise significant relationships following the Nemenyi post-hoc test, with * $p < 0.05$, ** $p < 0.01$, *** $p < 0.001$, **** $p < 0.0001$.* ***d*** *Networks coarse-grained after MIND estimation had reduced correlations with gene co-expression. Lines show the mean and shaded areas show ±1 standard deviation of correlation values across all non-empty combinations of donors used to estimate gene co-expression networks.*

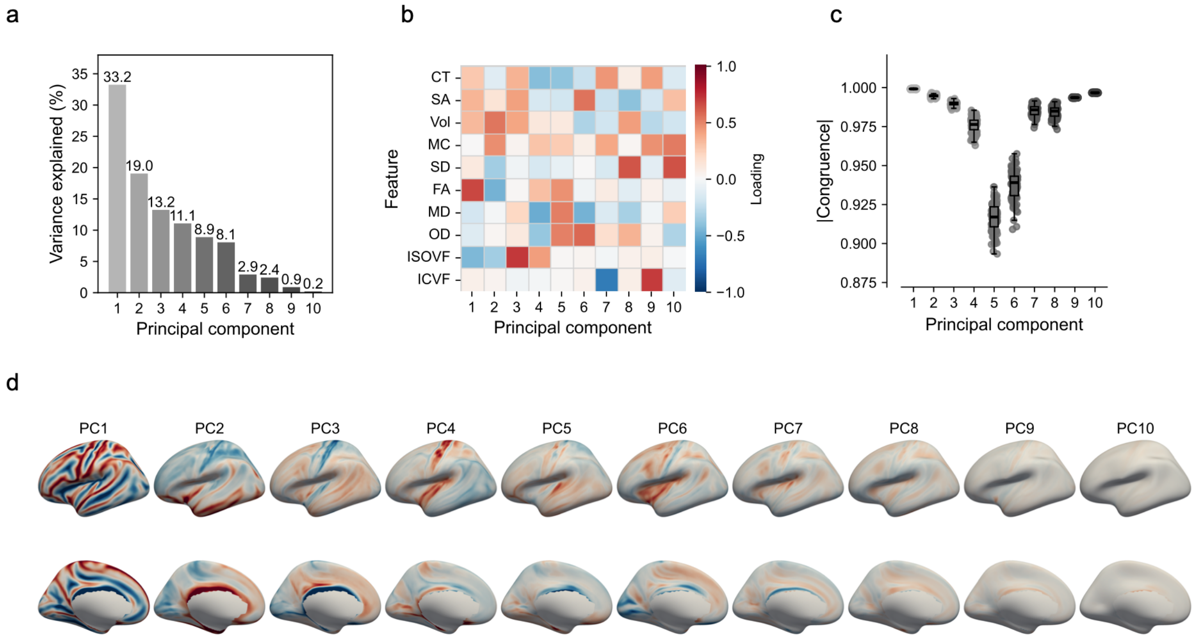


***Supplementary Figure 10: Principal component analysis of 10 macro- and microstructural features.*** ***a*** *Scree plot showing the variance explained by all 10 principal components.* ***b*** *Matrix showing the loadings of each MRI-derived morphological feature on the 10 principal components.* ***c*** *Bar plots showing the absolute congruence between principal components computed on the full dataset and 100 random subsamples of N = 250 individuals. Except for PC5 and 6, the absolute congruence of loadings on all components and subsamples with the loadings computed on the full dataset is greater than 0.95, indicating that these components can be considered equivalent, and therefore stable, across subsamples. The slightly reduced congruences of PC5 and 6 may result from their explaining similar proportions of variance in the data, allowing them to arbitrarily switch positions in random subsets of the data.* ***d*** *Left hemisphere maps showing the vertex-wise loadings of each principal component obtained when projecting group mean morphological maps to component space.*

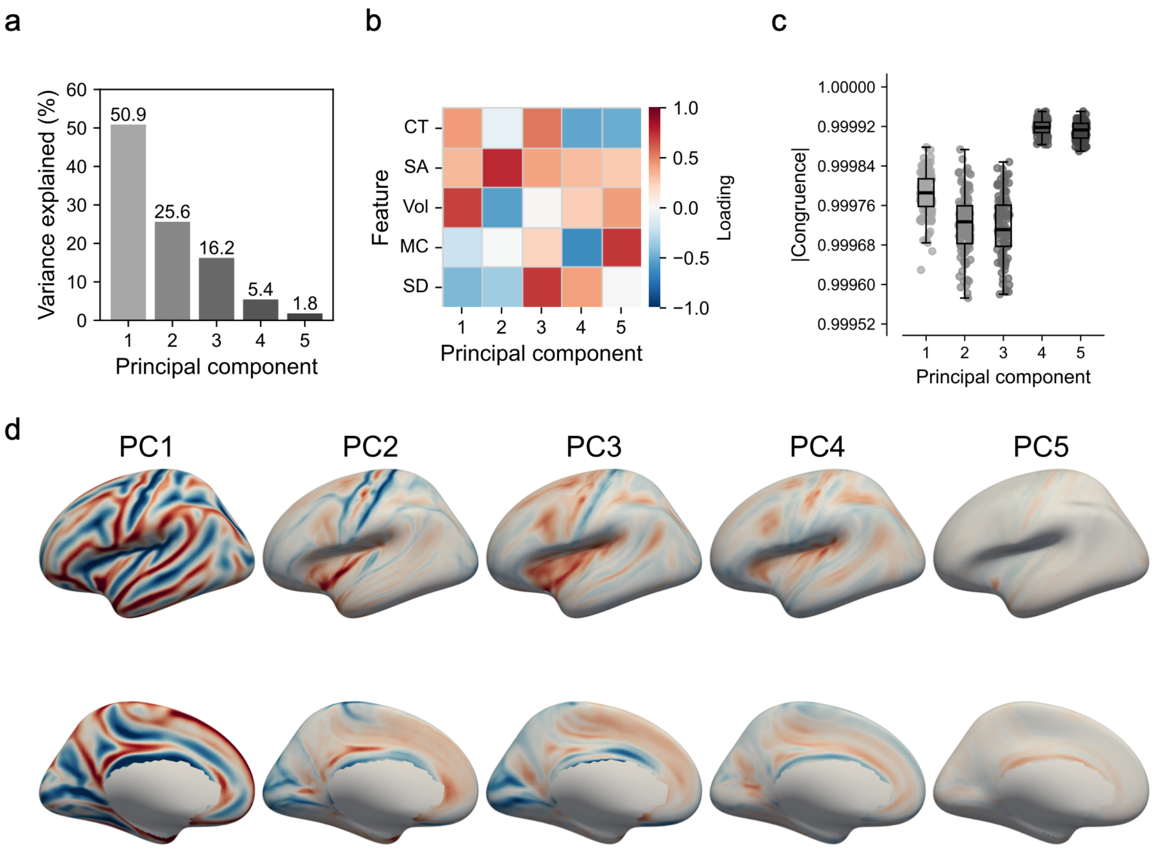


***Supplementary Figure 11: Principal component analysis of 5 macrostructural features. a*** *Scree plot showing the variance explained of all 5 principal components.* ***b*** *Matrix showing the loadings of each MRI-derived morphological feature on the 5 principal components.* ***c*** *Bar plots showing the absolute congruence between principal components computed on the full dataset and 100 random subsamples of N = 250 individuals. For all components and subsamples, absolute congruence is greater than 0.95, indicating that components can be considered equivalent, and therefore stable, across subsamples.* ***d*** *Left hemisphere maps showing the vertex-wise loadings of each principal component obtained when projecting group mean morphological maps to component space.*

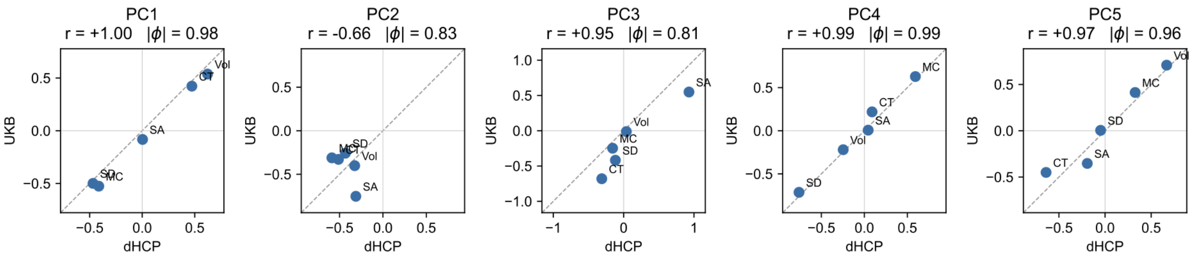


***Supplementary Figure 12: Feature loadings on principal components are externally replicable.*** *Scatterplots show the relationships between loadings of each macrostructural MRI feature on each of the five principal components derived separately in UKB and dHCP data. Pearson r correlation values and absolute congruences are displayed. Absolute congruence, |Φ|, is greater than 0.95 for PC1, 4, and 5, suggesting that these can be considered equivalent components, but not for PCs 2 and 3. This suggests that some large-scale and fine-scale patterns of cortical structural variation may be preserved even across drastically different age groups (adult vs. neonatal), acquisition parameters, and preprocessing pipelines.*

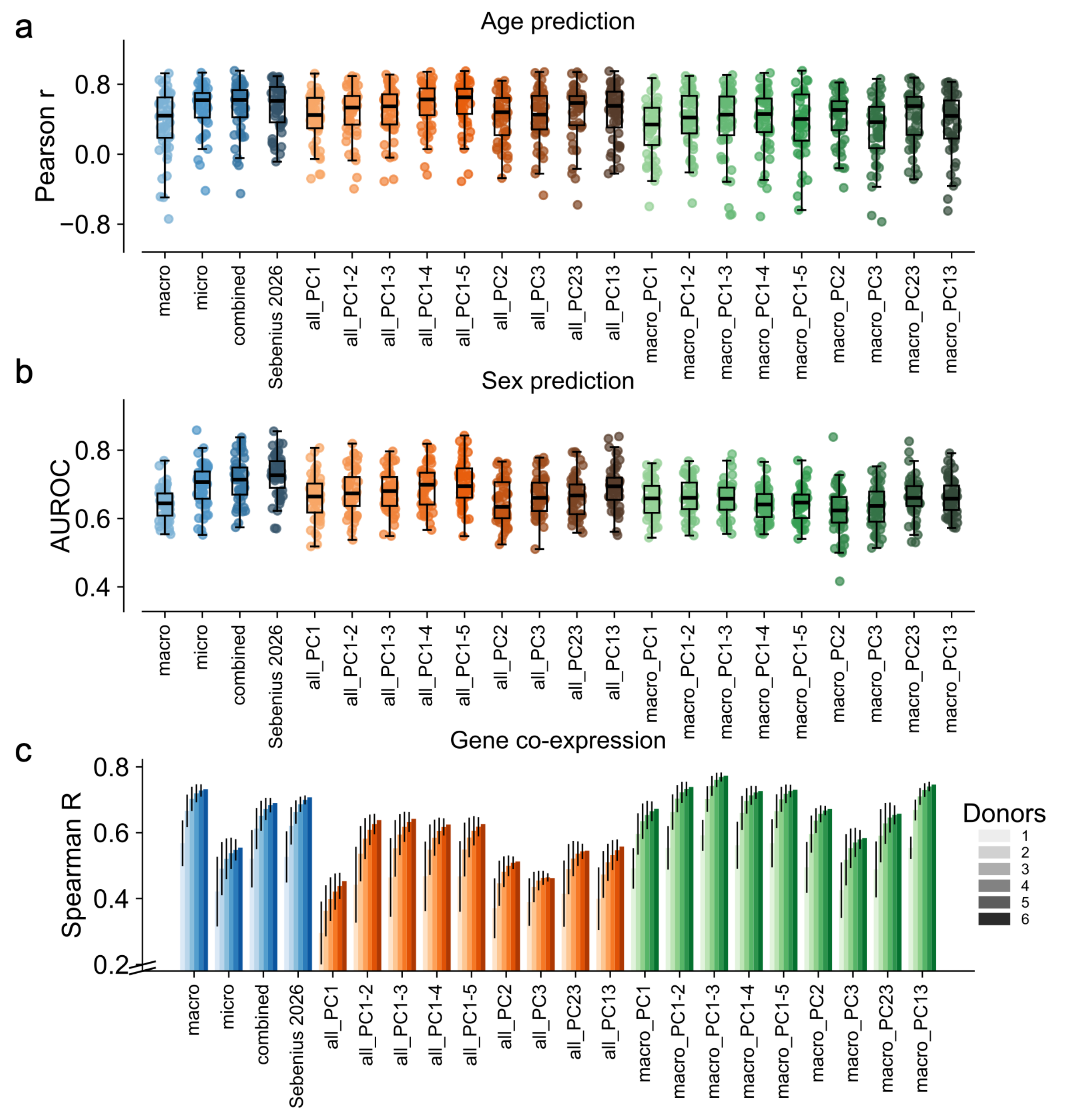


***Supplementary Figure 13: Biological validation of all feature sets.** In the following plots, blue colours signify MIND networks estimated using combinations of raw features, orange colours signify MIND networks estimated using combinations of principal components computed on all 10 macro- and microstructural features (‘all’), and green colours signify MIND networks estimated using combinations of principal components computed on 5 macrostructural features only (‘macro’). **a** Age prediction results. **b** Sex classification results. In **a-b**, pairwise significance bars are not shown due to space constraints. **c** Gene co-expression correlations. Bars show the mean ±1 standard deviation of correlation values across all non-empty combinations of donors used to estimate gene co-expression networks.*
